# Generation and characterization of conditional yeast mutants affecting each of the two essential functions of the scaffolding proteins Boi1/2 and Bem1

**DOI:** 10.1101/2022.06.15.496310

**Authors:** Abigail Sulpizio, Lancelot Herpin, Robert Gingras, Wenyu Liu, Anthony Bretscher

## Abstract

Boi1 and Boi2 are closely related yeast scaffolding proteins, either of which can perform an essential function. Previous studies have suggested a role in cell polarity, interacting with lipids, components of the late secretory pathway, and actin nucleators. We report detailed studies of their localization, dynamics, and the generation and characterization of conditional mutants. Boi1/2 are present on the plasma membrane in dynamic patches, then at the bud neck during cytokinesis. These distributions are unaffected by perturbation of the actin cytoskeleton or the secretory pathway. We identify two critical aromatic residues, present in both Boi1 and Boi2, in the essential C-terminal PH domain, that cause temperature sensitive growth resulting in defects in polarized growth leading to cell lysis. The scaffolding protein, Bem1, colocalizes with Boi1 in patches at the growing bud, and at the bud neck, the latter requiring the N-terminal SH3 domain of Boi1p. Loss of function of Boi1-SH3 domain renders Bem1 essential which can be fully replaced by a fusion of the SH3b and PB1 domains of Bem1. Thus, the two essential functions of the Boi1/2/Bem1 proteins can be satisfied by Bem1-SH3b-PB1 and Boi1-PH. Generation and characterization of conditional mutations in the essential function of Bem1 reveal a slow onset of defects in polarized growth, which is difficult to define a specific initial defect. This study provides more details into the functions of Boi1/2 and their relationship with Bem1, and presents the generation of conditional mutants that will be useful for future genetic analysis.

## Introduction

The mechanism by which cells polarize is a fundamental question in cell biology that has been especially well explored in the budding yeast (Chiou et al., 2017). For example, to generate a bud, the cell has to select an appropriate site, build a polarized actin cytoskeleton, and deliver post-Golgi vesicles to sites of growth. In order to coordinate the process, there are many essential components involved.

The first step in generating a new bud is to select a site to which cell growth will occur. This involves the local activation of the small RhoGTPase, Cdc42 that is activated by its GEF, Cdc24 (Bi & Park, 2012; Pruyne & Bretscher, 2000a, 2000b; Zhao et al., 1995; Ziman & Johnson, 1994). Active Cdc42 has many roles, initially directing the nucleation and assembly of actin cables through the formin protein Bni as well as participating in the assembly of septin proteins at the bud neck (Adamo et al., 2001; Brown et al., 1997; Chen et al., 1997; Evangelista et al., 1997; Zhang et al., 2008). The actin cables are then used as the tracks for the myosin-V Myo2 to deliver post-Golgi secretory vesicles, marked by the Rab Sec4, to sites of exocytosis (Kabcenell et al., 1990; Pruyne et al., 1998; Salminen & Novick, 1987). During transport, the secretory vesicles acquire the exocyst, a multi-protein complex that facilitates the tethering of secretory vesicles to the plasma membrane and is essential for fusion with the plasma membrane in a process also involving Cdc42 (Gingras et al., 2022; TerBush et al., 1996; Wu et al., 2010).

Considerable interest has focused on the initial single-site activation of Cdc42. Extensive studies have shown that this involves a local positive feed-back loop coordinated by the scaffolding protein Bem1 (Bose et al., 2001; Grinhagens et al., 2020; Leeuw et al., 1995; Liao et al., 2013; Park et al., 1997; Takaku et al., 2010; Yamaguchi et al., 2007). Bem1 is a modular protein containing two SH3 domains (SH3a and SH3b), a PX domain, and a C-terminal PB1 domain. A two-hybrid screen for interactors of Bem1’s second SH3 domain yielded a protein called Boi1 (Bem One Interactor 1), and its homolog, Boi2. Loss of either Boi1 or Boi2 has no effect on cell growth, whereas loss of both is lethal or highly deleterious (Bender et al., 1996; Matsui et al., 1996). Boi1 and Boi2 are large modular scaffolding proteins consisting of an SH3 domain, a SAM domain, followed by a proline-rich region that binds the Bem1SH3b domain and terminating in a Pleckstrin-Homology (PH) domain. Remarkably, the essential function of the Boi1/2 proteins can be supplied by just the Boi1 or Boi2 PH domain that is reported to bind Cdc42 (Bender et al., 1996; Kustermann et al., 2017; Matsui et al., 1996). Studies have shown that when the Boi proteins are depleted from the cell, secretory vesicles accumulate within the bud, and that the N-terminal Boi1 domain binds several components of the exocyst, and the C-terminal PH domain binds Sec1 (Kustermann et al., 2017; Masgrau et al., 2017). It was also found that the cells lacking Boi1/2 were capable of survival with a gain-of-function mutant of the exocyst complex, *EXO70\** (Masgrau et al., 2017). The collective results suggest that Boi1/2 proteins coordinate cell polarization with secretory vesicle exocytosis. Additionally, Boi1/2 have been reported to bind through their N-terminal region directly to the formin Bni1 and accessory protein Bud6 that are involved in the nucleation of actin filaments (Glomb et al., 2020).

Early work has also shown that an inactivating mutation in the SH3 domain of Boi1 in *boi2*Δ cells (*boi1W53K boi2*Δ) renders *BEM1* essential (Bender et al., 1996). Thus, the Boi1/2 proteins have one essential function requiring the PH domain, and a second essential function overlapping with Bem1. In our previous work, we have made extensive use of fast-acting conditional mutants to *TPM1, MYO2* and *BNI1* that have been very useful in pinpointing their essential functions (Chernyakov et al., 2013; Evangelista et al., 1997; Pruyne & Bretscher, 2000b; Pruyne et al., 1998; Schott et al., 1999). In this study, we set out to examine the distribution of Boi1/2 and Bem1 in more detail and generate conditional mutations in each of the two essential functions as an unbiased approach to explore their precise roles.

## Results

### Boi1/Boi2 are essential proteins localized to short-lived polarized patches on the plasma membrane

In order to study the cellular functions of Boi1/2, we confirmed that deletion of either of the two genes confers no apparent growth defect, yet loss of both is lethal indicating that Boi1 and Boi2 share an essential function in our genetic background (BY4741/2) (Supplemental 1.1 and 1.2). Previously, Boi1 and Boi2 have been shown to be generally polarized throughout the cell cycle and at the bud neck during later growth (Hallett et al., 2002; Kustermann et al., 2017; Masgrau et al., 2017). Here, we localize Boi1 and Boi2 C-terminally tagged endogenously with mNeonGreen (mNG) at a higher spatial and temporal resolution than prior studies. During polarized growth, both proteins localized to the nascent bud and in patches associated with the cell cortex. (Figure 1A and B). We confirmed that this localization was only at the plasma membrane, and not on internal structures, as single confocal slices through the middle of the cell exclusively show cortical association (Figure 1C and D). During late bud growth, the proteins shift localization to the bud neck. By imaging alongside the class-II myosin, Myo1, tagged with mScarlet, Boi1 was found to localize at the bud neck for 7-8 mins before completion of cytokinesis marked by the loss of Myo1-mScarlet and then remains there (Figure 1E)(Bi et al., 1998; Okada et al., 2021).

**Figure 1:**
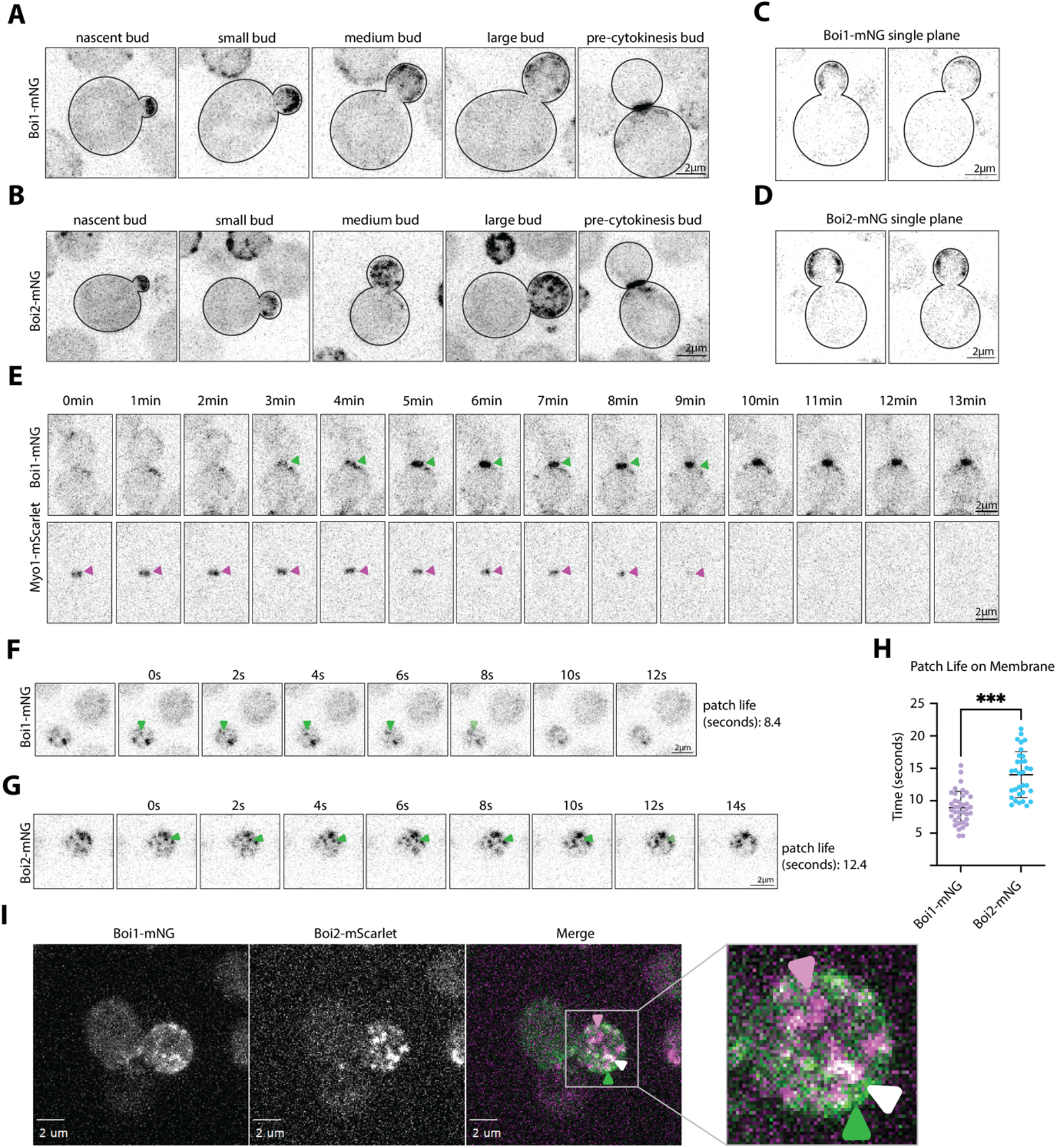
Boi1-mNG and Boi2-mNG localize to short-lived stable polarized patches on the plasma membrane and at the bud neck during late bud growth. (A and B) Images of endogenously tagged Boi1 or Boi2 with mNeonGreen taken at five stages of cell growth. Captures were at 200ms exposure with max-projection of 15 stacks covering 4.2µm of the cell in the z-plane. (C and D) Single slices of Boi1-mNG and Boi2-mNG showing localization is only at the plasma membrane and not on cytoplasmic vesicles. Images of Boi1-mNG with Myo1-mScarlet. Images are z-projections of 15 planes covering 4µm of the cell. Arbitrary timepoint “0min” is 3 minutes before arrival of Boi1-mNG to the bud neck. Figures are representative of at least 5 examples. Scale bars are 2µm. (F and G) Stills of Video 1 and 2. Single-plane timelapse of Boi1-mNG or Boi2-mNG captured every 160ms with 50ms exposure. (H) Graph showing patch lifetimes of Boi1-mNG and Boi2-mNG in the growing cell. The mean patch longevity for Boi1-mNG is 9 seconds, and Boi2-mNG is 14 seconds. Timing of patch longevity is significantly different p < 0.001 through a Mann-Whitney statistical test. (I) Co-localization of Boi1-mNG and Boi2-mScarlet, max-projections of 15 stacks covering 4.2µm of the cell. Boi1-mNG is exposed for 100ms, and Boi2-mScarlet for 300ms. White arrows indicate instances of colocalization, green Boi1-mNG alone, and magenta Boi2-mScarlet alone. All scale bars are 2µm.

To assess relative dynamics of Boi1/2 patches on the membrane, the lifetime of single patches was measured (Figure 1F and G, Video 1 and 2). Boi1 patches were found to be shorter-lived with a mean lifetime of about 9s, whereas Boi2 patches persisted longer, with a lifetime of about 14s, a timing consistent with earlier measurements (Figure 1H) (Gingras et al., 2022). To explore whether Boi1 and Boi2 patches colocalize, cells were imaged expressing Boi1-mNG together with Boi2-mScarlet. Although both proteins colocalized, there was not perfect overlap with clear instances of Boi1 and Boi2 localizing to separate patches (Figure 1I, example video: Video 3). This lack of colocalization is not due to the increased abundance of Boi2-mScarlet or the longer lifetime since Boi1-mNG is seen to be in independent patches lacking Boi2-mScarlet. Thus, Boi1 and Boi2 are similarly localized to short-lived patches that presumably participate in their overlapping essential function, yet each may also play additional distinct, subtle, roles.

### Boi1 puncta are more dynamic than Boi2 puncta

As the Boi1-mNG patch life on the membrane is shorter than Boi2-mNG, we next examined the protein turnover within single patches using Fluorescence Recovery After Photobleaching (FRAP). Individual cortical patches were bleached and recovery was monitored for Boi1-mNG and Boi2-mNG (Figure 2A, B, C, and D). Boi1-mNG fluorescence appeared to recover much faster than Boi2-mNG, with most of the Boi2-mNG patches never recovering after photobleaching (Figure 2E). This suggests that Boi1’s shorter patch lifetime may reflect a higher protein turnover relative to Boi2.

**Figure 2:**
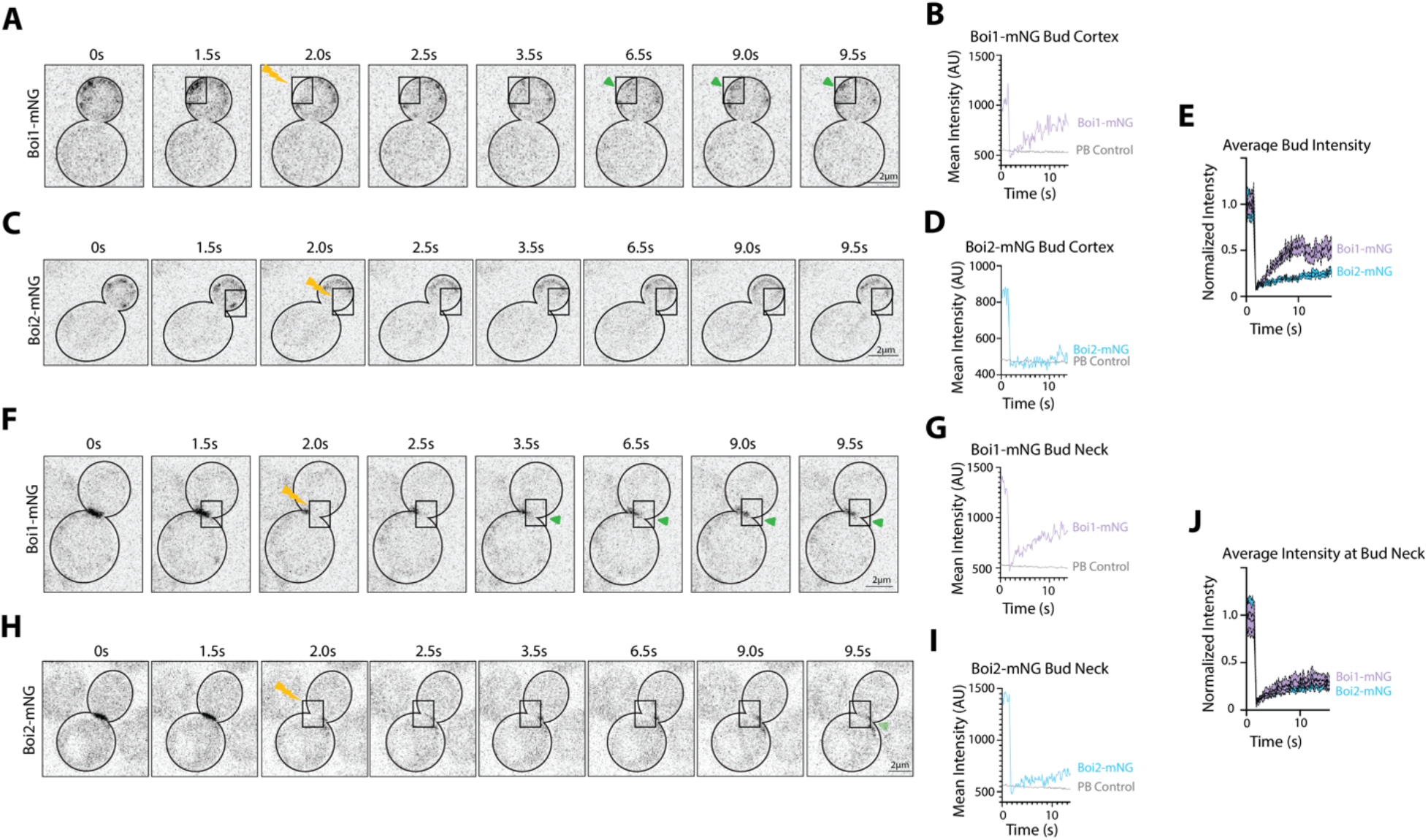
Boi1 is more dynamic than Boi2 at patches within the bud and at the bud neck. (A) Boi1-mNG Fluorescence Recovery After Photobleaching (FRAP) images before, during, and seconds after photobleaching (depicted with yellow lightning bolt) a region in the growing bud (outlined with box). Images are single z-plane slices in the middle of the cell at a 50ms exposure. Timelapse was taken with 160ms between each frame. (B) Trace of the intensity of the patch region compared to the photobleaching control (PB Control) of another mother cell in the field of view. (C) Boi2-mNG FRAP images of a mid-budding cell taken with the same parameters. (D) Trace of intensity of that region compared to the photobleaching control. (E) Average normalized intensity of Boi1-mNG (lilac, n=10) and Boi2-mNG (blue, n=6) traces within the bud overlaid with one another. (F and H) FRAP imaging of Boi1-mNG and Boi2-mNG at the bud neck taken with the same parameters as A and C with photobleaching of half the signal at the neck. (G and I). Subsequent traces of intensity. (J) Average normalized intensity of traces of Boi1-mNG (n=4) and Boi2-mNG (n=8) at the bud neck. Images were normalized to the highest average intensity. (K) Examples of laterally mobility in Boi1-mNG and Boi2-mNG patches on the cortex. Arrows indicate a specific patch that moves from the first timepoint on the membrane. All scale bars are 2µm.

To assess the dynamics of Boi1 and Boi2 at the bud neck, half of the signal at the bud neck was bleached and fluorescence recovery recorded (Figure 2F, G, H, and I). While Boi1 fluorescence appeared to recover more readily than Boi2, neither protein recovered particularly well (Figure 2J). Furthermore, patches hardly move laterally on the cortex (Supplemental 2.1). These experiments illustrate that Boi1 is more dynamic than Boi2 in both protein turnover at the patches and slightly at the bud neck. Boi1 is also more dynamic in patch longevity, highlighting a difference between the two proteins’ dynamics.

### Boi1-mNG and Boi2-mNG’s localization is independent of actin and secretory vesicle transport

In order to examine the relationship between Boi1 and Boi2 patches and other polarized structures, we initially assessed their localization relative to actin. Previous studies have reported that Boi1, and the *S. pombe* homolog Pob1, are capable of re-directing actin filament formation and interacting with actin-nucleator formins (Glomb et al., 2020; Rincón et al., 2009; Toya et al., 1999). Neither Boi1-mNG nor Boi2-mNG colocalize with actin cortical patches (Figure 3A and B) indicating they are not associated with the endocytic machinery. Consistent with this lack of colocalization, treatment of cells for five minutes with the Arp2/3 inhibitor CK-666, which results in the complete disassembly of actin patches (Hetrick et al., 2013; Nolen et al., 2009), has no impact on the polarized localization of Boi1 or Boi2 (Figure 3C).

**Figure 3:**
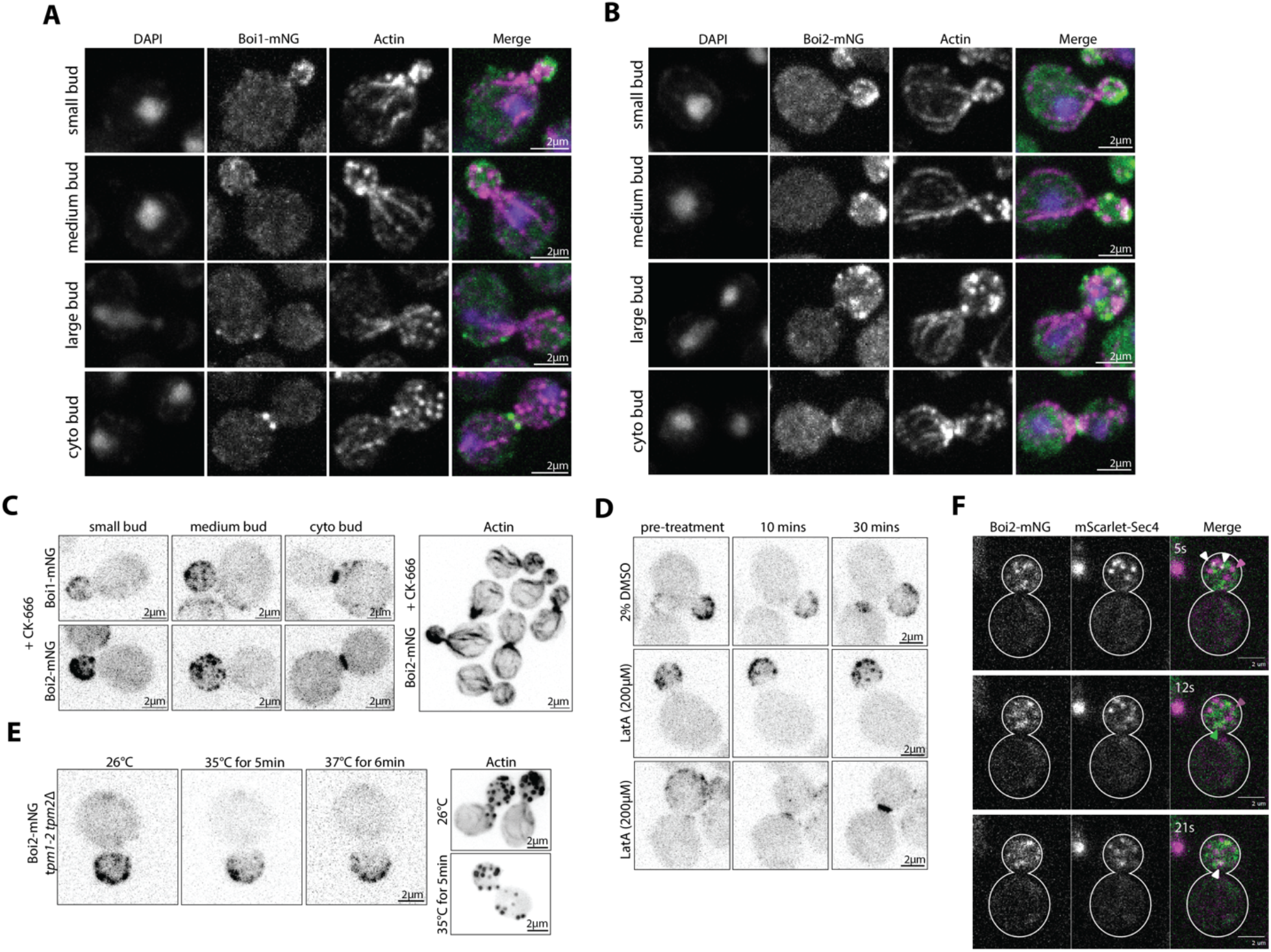
Boi1-mNG and Boi2-mNG localization on the plasma membrane and at the bud neck is independent of actin cortical patches, cables, and secretory vesicles. (A) Fixed cell imaging of Boi1-mNG (200ms exposure), phalloidin (200ms exposure), and DAPI staining (100ms exposure) at four different stages during budding, small bud, medium, large, and during cytokinesis. All images are maximum z-stacks of 15 planes covering ∼4.2µm. (B) Images of Boi2-mNG in four different stages of the cell cycle with the same parameters as in A. (C) Boi1-mNG and Boi2-mNG after CK-666 treatment, images taken within 10-20 minutes after treatment. Both images are maximum z-stacks of 15 planes covering ∼4.2µm. Exposure 200ms for both cell types. Second panel is actin phalloidin staining (75ms exposure) fixed after 5 minutes of CK-666 treatment. (D) Boi2-mNG with either treatment with a 2% DMSO control or Latrunculin A (200µM) two examples of LatA treatment show localization during mid and late bud cycle. Images were taken with 100ms exposure, and max-projections of 15 planes covering ∼4.2µm. (E) Boi2-mNG in *tpm1-2 tpm2Δ* cells. Max projections of 15planes ∼4.2µm. Exposure at 100ms. Second panel is actin phalloidin staining at 25ms exposure. (F) Boi2-mNG and mScarlet-Sec4 colocalization. 50ms exposure for each. Images taken in a single plane at the bottom of the cell. Arrows indicate individual Boi2-mNG localization (green), individual mScarlet-Sec4 localization (magenta), or co-localization (white). See Video 4 for more detail on mScarlet-Sec4 localization compared to Boi2-mNG. All scale bars are 2µm.

To determine if Boi1/2 patches are dependent on actin cables, we examined Boi2-mNG localization in the *tpm1-2 tpm2*Δ mutant which rapidly disassembles cables upon shift to the restrictive temperature (Pruyne et al., 1998). After shifting *tpm1-2 tpm2*Δ cells expressing Boi2-mNG to 35°C for 5 minutes, Boi2 polarization remained unchanged (Figure 3E), indicating that the polarization of Boi2 patches is independent of the presence of actin cables. Consistent with this, treatment of cells for 10 minutes with the drug Latrunculin-A, which results in disassembly of the entire actin cytoskeleton (Ayscough et al., 1997; Spector et al., 1989), had no impact on the presence of polarized Boi2 patches at the cell cortex or the bud neck (Figure 3D). Thus, the localization of Boi2 is independent of the actin cytoskeleton, consistent with earlier observations (Kustermann et al., 2017).

Previous studies have implicated Boi1/2 in exocytosis, showing that after depletion of Boi1/2, the cell accumulates secretory vesicles (Kustermann et al., 2017; Masgrau et al., 2017). To examine the relationship of Boi2 to secretory vesicles, we co-imaged Boi2-mNG with the secretory vesicle marker, mScarlet-Sec4. Strikingly, the majority of Boi2-mNG patches do not colocalize with secretory vesicles, or vice versa, although some colocalization of tethered vesicles and Boi2 patches does occur (Figure 3F, Video 4). The majority of non-colocalization is at least partially due to the relatively short tether-to-fusion time of secretory vesicles and the number of untethered vesicles in the bud (Gingras et al., 2022). Furthermore, examination of videos shows that secretory vesicles tether in places apparently devoid of Boi2-mNG patches. This led us to question whether these vesicles are tethering with regions which have no Boi1/2 present, or if they’re merely tethering at Boi1 patches. To answer this, we looked at Boi2-mNG mScarlet-Sec4 in a *boi1*Δ. In this cell type, the mScarlet-Sec4 vesicles had a similar localization and movement as the cells in an otherwise wildtype background (Video 5), indicating that mScarlet-Sec4 vesicles are capable of tethering on the membrane at sites lacking Boi1/2.

Given their reported role in exocytosis of secretory vesicles, we examined whether the localization of Boi1 or Boi2 were influenced in mutants with conditional defects in the late secretory pathway. Localization of Boi1-mNG or Boi2-mNG to polarized patches is unaffected by shifting the conditional *sec4-8* mutant to the restrictive temperature for 30 minutes (Supplemental 3.1), a condition that compromises both vesicle transport and overall secretion (Harsay & Bretscher, 1995; Salminen & Novick, 1987). Thus, the polarized patch localization of Boi1 and Boi2 is independent of polarized secretory vesicle delivery and the regulation of this localization is upstream of exocytosis.

Earlier studies have reported that Boi2 is localized to the cytosol in the conditional exocytic mutant *sec6-8* after shifting to the restrictive temperature for 1-2 hours (Masgrau et al., 2017). To explore this phenomenon, we shifted the conditional exocytic mutant *sec6-4* to the restrictive temperature for 30 minutes. Unlike the *sec4-8* conditional mutant, the *sec6-4* mutation only compromises vesicle tethering, not transport, at the restrictive temperature. Interestingly, even prior to shifting, Boi1-mNG appeared to localize to depolarized cortical patches in the mother – a localization never observed in wildtype cells – which became more prominent at the restrictive temperature (Supplemental 3.2 and 3.3). Together, these results indicate that the presence of Boi1/2 in polarized cortical patches is not directly influenced by the delivery of secretory vesicles nor a requirement for exocytosis; however, patch regulation may be influenced by exocyst function.

### Boi1PH is responsible for Boi1 cortical bud localization while Boi1SH3 is important for Boi1 and Bem1 localization at the bud neck

After our initial characterization of Boi1/2, we then wanted to understand the contributions of the individual domains. It has previously been shown that the Boi1/2 Pleckstrin-Homology (PH) domain including the C-terminal tail represents the minimal essential domain which facilitates interactions with both the plasma membrane and Cdc42. Boi1 and Boi2 also each contain an SH3 domain, a SAM motif, and a proline-rich region necessary for interaction with Bem1 (Figure 4A) (Bender et al., 1996; Hallett et al., 2002; Kustermann et al., 2017; Matsui et al., 1996). We focused on the PH and SH3 domains by first replacing expression of the endogenous *BOI1* with just the Boi1 PH domain (*boi1(730-980))*, in a *boi2*Δ background to generate *Boi1PH boi2*Δ. This strain grew normally at all temperatures (Figure 4B), confirming that just the PH domain can provide the Boi1/2 essential function. Upon tagging Boi1PH with the mNeonGreen, the patch-like distribution is diminished and Boi1PH-mNG is only polarized to the bud and not found at the bud neck (Figure 4C). Examining a few optical sections revealed that Boi1PH-mNG is restricted to the cortex and not found on internal structures (Figure 4D, Video 6). These results confirm that the shared essential function mediated by the PH domain is at the cell cortex, and another part of the protein is responsible for the patch-like distribution that the wildtype-protein exhibits. It has been suggested by previous work that the protein’s SAM domain might help facilitate homodimerization (Jia et al., 2020). This may explain why the patch-formation is eliminated in the Boi1PH-mNG cells.

**Figure 4:**
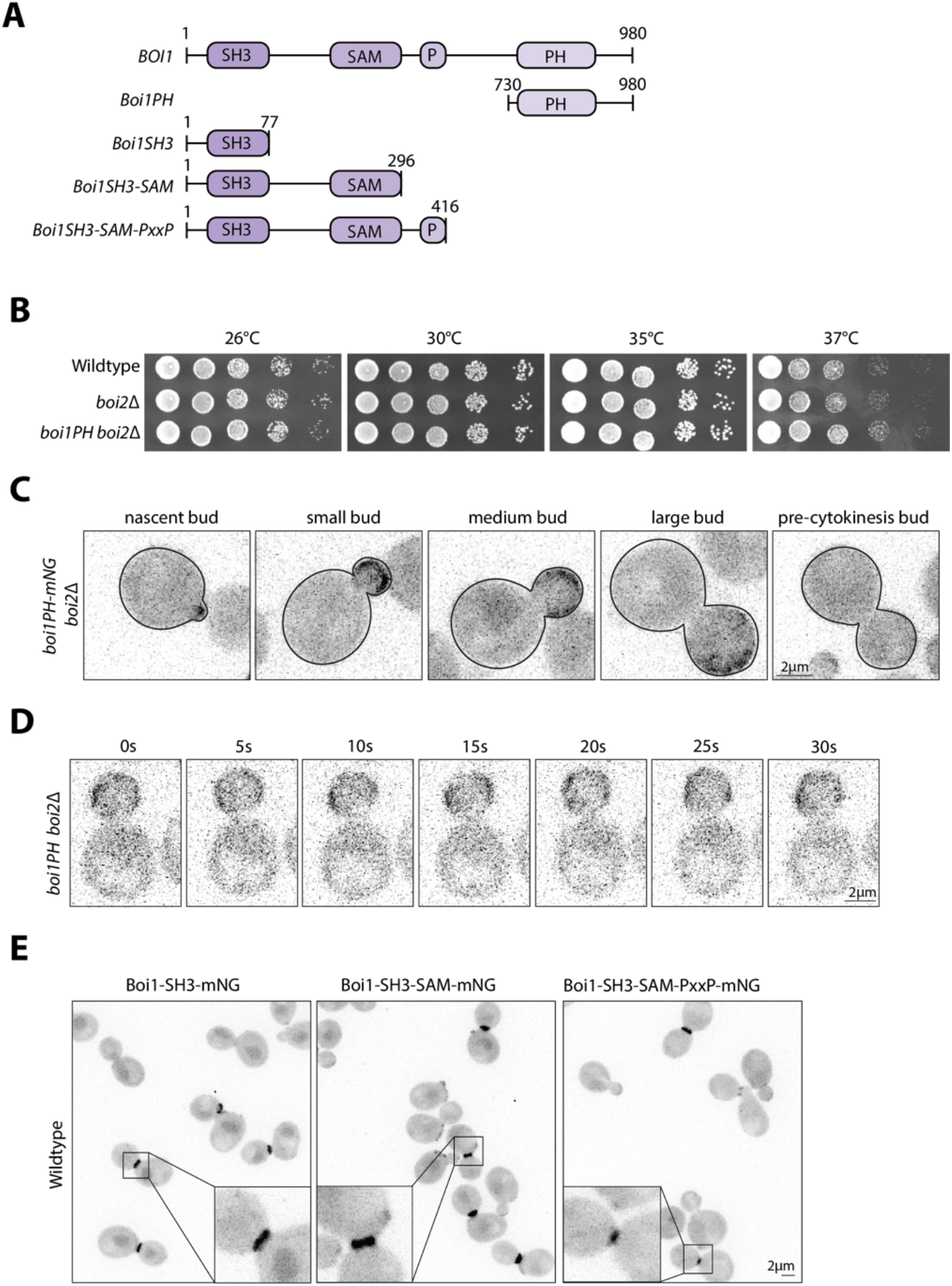
Boi1PH is the essential domain important for stable Boi1 cortical membrane localization, and the Boi1SH3 domain is important for Boi1 and Bem1’s localization at the neck. (A) Schematic showing the truncations of BOI1 used in this figure. (B) Dilution assays of wildtype, *boi2Δ*, and *boi1PH(730-980) boi2Δ* cells at 26°C, 30°C, 35°C, and 37°C all on YPD plates grown for 2 days. (C) Max projections of 15 z-planes of *boi1PH-mNG boi2Δ* cells at the nascent, small bud, medium bud, large, and pre-cytokinetic bud. Images were exposed for 200ms. Images cover ∼4.2µm of the cell. (D) Stills of Video 6 showing *boi1PH-mNG boi2Δ* cells. Images were taken with 50ms exposure max projections of 6-planes at the bottom of the cell. (E) Images of three different tagged truncations of Boi1 in a wildtype cell. Images were taken at 500ms exposure, max projections with 13 z-planes covering ∼4µm. All scale bars are 2µm.

After initial characterization of the Boi1PH domain, we explored the localization of the N-terminal part of the protein. Tagged versions of three different truncations of Boi1 (Boi1SH3, Boi1SH3-SAM, Boi1SH3-SAM-PxxP) all localized exclusively to the bud neck and were never found at the bud cortex (Figure 4E). Thus, the cortical bud localization of Boi1 is mediated by the PH domain, whereas the bud neck localization is mediated by the Boi1 SH3 domain.

### Generation of temperature sensitive mutations in Boi1 highlight two aromatic residues in the PH domain

To investigate the essential function of Boi1/2 further, we used PCR mutagenesis to generate temperature sensitive alleles of *BOI1* in a *boi2*Δ background. Two different strategies were employed to incorporate the mutated gene. The first was direct integration of the mutated gene into the genome using an integrating plasmid, and the second employed a plasmid shuffle screen (Figure 5A). Using each strategy in three separate screens for mutants that could grow at 26°C but not at 35°C, we were unable to recover conditional mutants using the integration approach, but recovered several from the plasmid shuffle screens. Three alleles, *boi1-1, boi1-2*, and *boi1-3*, were selected for further study. Interestingly, they failed to grow at 35°C, but growth was restored at 37°C. All three mutants were recessive, as expression of the wildtype *BOI1* or *BOI2* genes suppressed their temperature sensitive phenotype (Figure 5B). Multiple attempts were made to introduce these mutations into the chromosome, but all failed. We conclude that they are lethal at the chromosomal locus, so all further studies were performed with them expressed from a centromeric plasmid.

**Figure 5:**
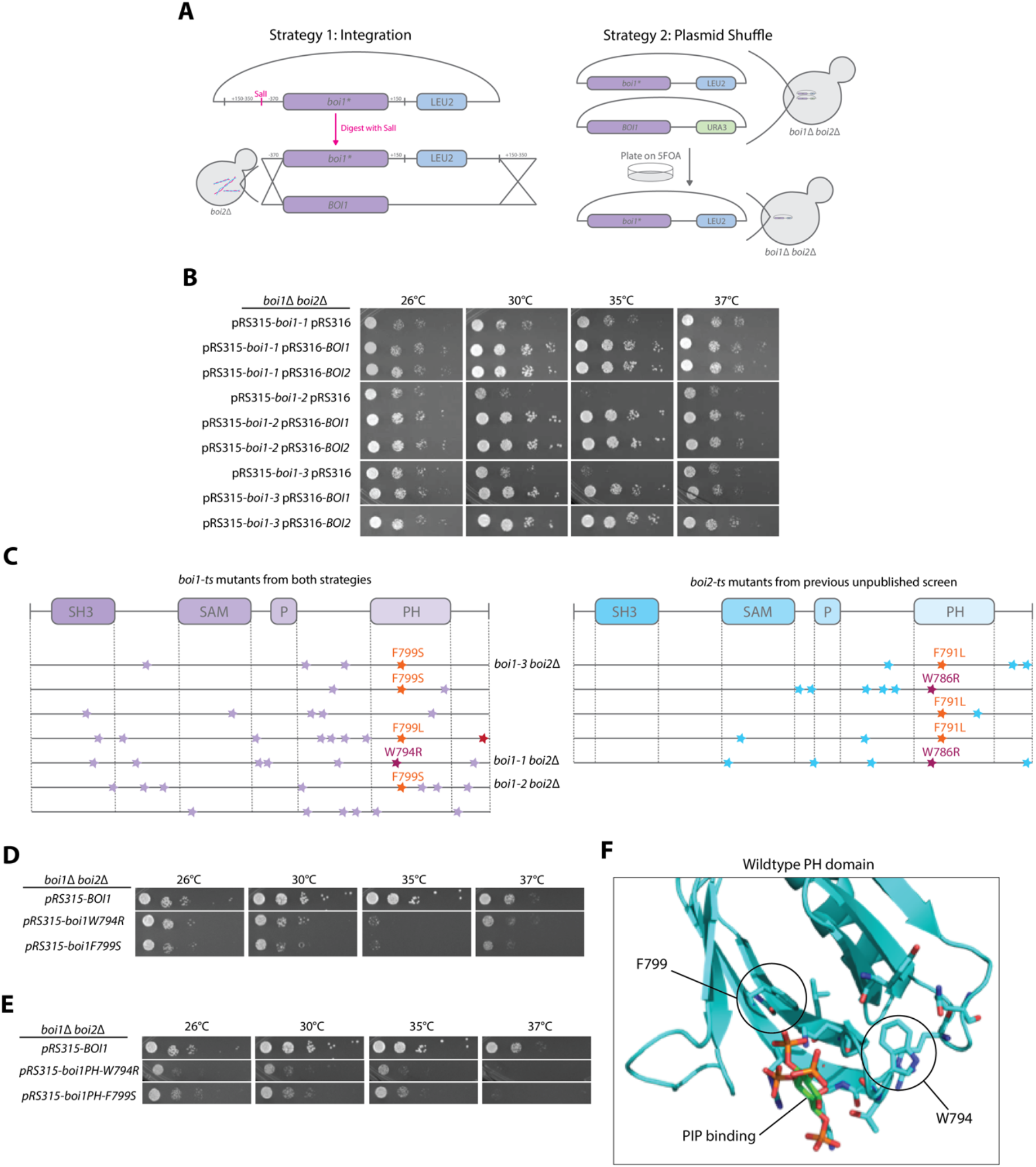
Generation of conditional mutants in *BOI1*. (A) Strategies of the screen to develop the temperature sensitive mutants in *BOI1* in a *boi2Δ* background. Strategy 1 highlights integration of the mutated gene to replace the endogenous gene, and Strategy 2 illustrates a plasmid shuffle assay in a *boi1Δ boi2Δ* background. Each screen was done 3+ times and about 15,000 clones were analyzed. (B) Growth of three *boi1* conditional mutants. Each mutant is recessive as the temperature sensitivity was suppressed by incorporation of either *BOI1* or *BOI2* on a centromeric plasmid. Dilution assays were conducted on SC -LEU/-URA plates and grown for 2 days at the respective temperatures. (C) Domain maps highlighting mutation variance across *BOI1* and *BOI2*, purple/blue stars indicate single nucleotide mutations changing the amino acid. Highlighted mutations show similarities across screens with F791/F799 and W786/W794 as the two recurrent mutations found in each candidate. (D) Single W794R or F799S mutations introduced into full length *BOI1* in *boi1Δ boi2Δ* cells show temperature sensitive growth. Cells were grown on SC -LEU media for 2 days at the respective temperatures. (E) Single W794R or F799S mutations introduced into *Boi1PH* in *boi1Δ boi2Δ* cells show temperature sensitive growth. Cells were grown on -LEU plates for 2 days. (F) Homology modeling shows where the two mutations potentially fall in the protein structure. Model was made with human DAPP1 PH domain bound to inositiol to represent PIP_2_ binding.

Interestingly, although multiple mutations were found in the temperature sensitive alleles, several of them had mutations in one of two aromatic residues in the PH domain of *BOI1*. Remarkably, in a previous unpublished screen for conditional mutations in *BOI2* in a *boi1*Δ background we again found the corresponding residues affected (Figure 5C, Supplemental 4.1). These mutations were either the W794R or the F799S in *BOI1* or W786R and F791S in *BOI2*. These individual mutations were found to confer temperature sensitive growth when individually placed in the full-length protein or the Boi1PH version (Figure 5D and E). Although single mutations caused temperature sensitivity, the double W794R F799S mutant was not viable (Supplemental 4.2). Both W794 and F799 are adjacent to the PIP_2_ binding pocket of Boi1, potentially explaining the temperature sensitivity (Figure 5F). To further explore what residues caused temperature sensitivity at Boi1 positions 794 and 799, we generated libraries of Boi1 with random codons at these two positions and asked which of these were temperature sensitive when introduced into a *boi1*Δ *boi2*Δ background. Among several colonies, we found: W794R, W794G, F799S, F799L, and F799G, (Supplemental 4.3). These results establish the critical and restrictive nature of residues FW794 and F799 in Boi1.

### The *boi1* conditional mutants exhibit defects in cell morphology, secretion, and actin patch localization

To better characterize which cellular process Boi1/2’s essential function supports, we observed the growth phenotypes of the *boi1-ts*. The *boi1* conditional mutants did not cease growth immediately upon shifting to the restrictive temperature (Supplemental 5.1) and showed variable viability after shifting to 35°C for 4h, with about 50% loss for *boi1-1* and *boi1-2*, and 100% for *boi1-3* (Supplemental 5.2). As *boi1-1* exhibits the least clear temperature sensitivity, our characterizations focused on *boi1-2* and *boi1-3* (both containing the F799S mutation).

To explore Boi1’s essential function, we compared the growth morphology of *boi1-2 boi2*Δ cells to well characterized mutants affecting either the actin cytoskeleton, or exocytosis. Upon shifting to 35°C, *boi1-2* cells showed a variable phenotype of diminished cell growth and then eventually lysing (Figure 6A). By contrast, *tpm1-2 tpm2*Δ cells, which lose actin cables at the restrictive temperature (Pruyne et al., 1998), exhibit altered growth and also eventually lyse (Figure 6B). Conditional defects in secretion as exhibited by a *sec4-8* mutant, simply halt cell growth at the restrictive temperature and do not lyse (Figure 6C). This phenotypic analysis shows that the essential role of Boi1 is not restricted to organizing actin, or to secretion, but has a more complex function. The mutant most like the *boi1-2* was the conditional *pkc1-2* mutant affecting the essential kinase activity of the cell wall integrity pathway (Levin et al., 1990; Paravicini et al., 1992). This mutant appears to have a similar lysing pattern which shows lysing after about 80 minutes at the restrictive temperature (Figure 6D).

**Figure 6:**
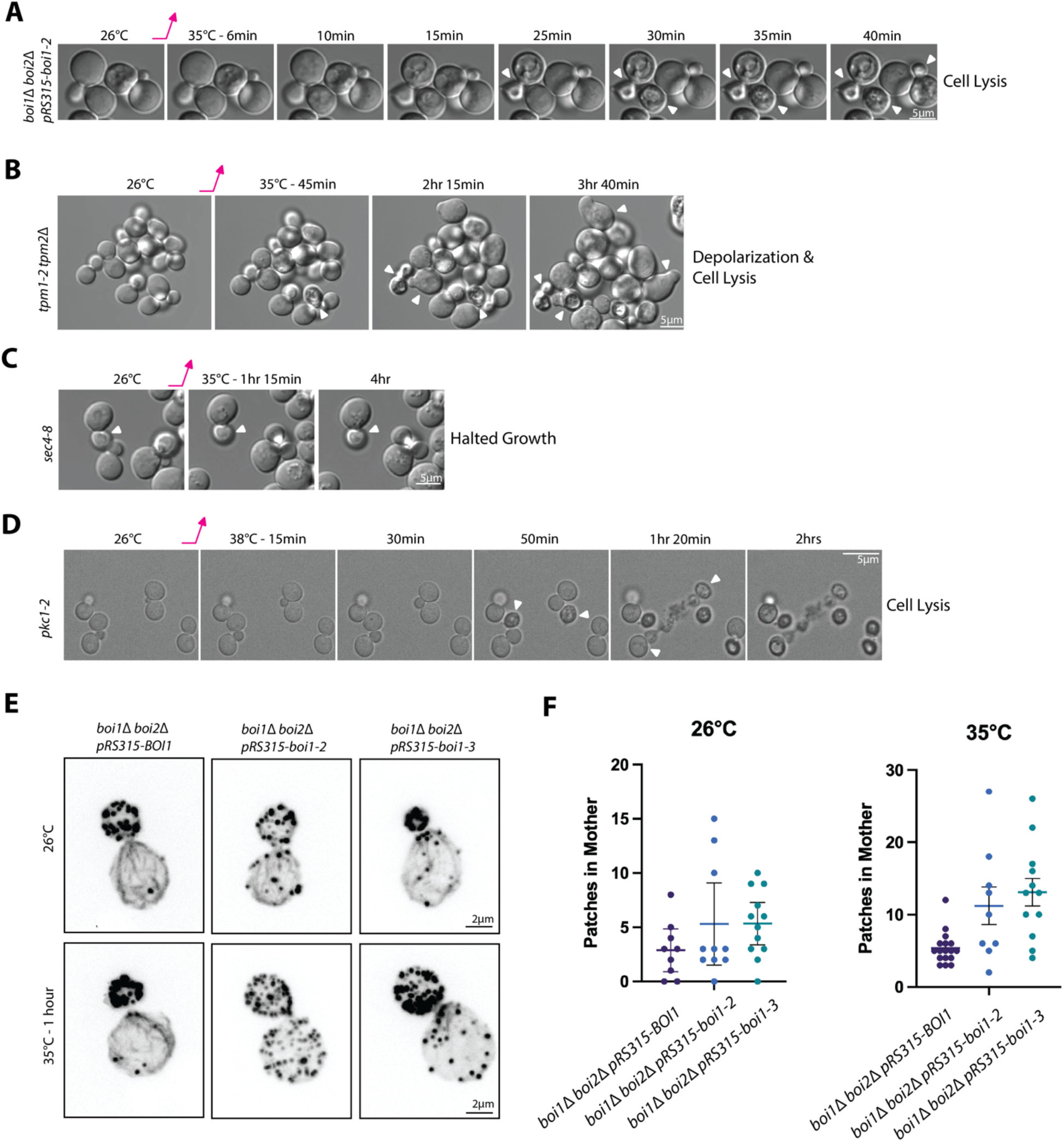
Characterization of the boi1 conditional mutants. (A) Morphology of *boi1*Δ *boi2*Δ pRS315-boi1-2 at 26°C and then shifted to 35°C for several minutes. Microscopy images are stills taken with DIC microscopy to look at overall cellular structure. Scale bars are 5µm. (B, C, and D) Morphology of different mutants related to secretion and cell wall integrity for comparison: *tpm1-2 tpm2Δ, sec4-8*, and *pkc1-2*, respectively. Images were taken with the same parameters as A. (E) Actin morphology of cells grown at 26°C and at 35°C for 1 hour and fixed with PFA. Cells were stained with phalloidin-568 (See methods) and microscopy is 15 z-stack planes with 100ms exposure. Scale bars are 2µm. (G) Graphs showing the number of patches in the mother in cells at 26°C and shifted to 35°C for 1 hour: 26°C BOI1 (n=9), boi1-2 (n=10), and boi1-3 (n=12); at 35°C BOI1 (n=17), boi1-2 (n=9), and boi1-3 (n=12).

The actin cytoskeleton was partially disrupted in the *boi1* conditional mutants. Whereas wildtype cells have cortical patches almost exclusively in the bud and robust actin cables whether grown at 26°C or shifted to 35°C for one hour, *boi1-2* cells lost much of the patch polarity even at 26°C with diminished cables, with even greater loss of patch polarity and cables at 35°C (Figure 6E). Similarly, *boi1-3* lost polarity of patches and had reduced cables after shifting to 35°C (Figure 6E). Quantitation of patch localization showed significant differences for patch polarity in both the *boi1-2* and *boi1-3* cells (Figure 6F). Our phenotypic characterization of the conditional *boi1* mutants did not reveal a clear role in either secretion or cytoskeletal polarity making its essential function difficult to pinpoint.

### *BOI1* participates in the Cdc42 pathway and the full-length is influenced by PIP_2_ concentrations

Earlier work has suggested that the Boi1/2 proteins function in the Cdc42 and potentially Rho3 pathways (Bender et al., 1996; Jia et al., 2020; Liao et al., 2013; Matsui et al., 1996). Cdc42 is an essential conserved small GTPase that regulates cell polarity from yeast to vertebrates (Chiou et al., 2017; Johnson, 1999) and its GEF, Cdc24 is also essential in yeast. Over-expression of either *CDC42* or *CDC24* on a 2µ vector suppressed the growth defect of *boi1-3* (Supplemental 6.1, Panel A). To see how specific this suppression was to Cdc42, each of yeast’s six small GTPases of the Rho family were expressed from 2µ vectors. *CDC42* suppressed well, with only *RHO3* over-expression also showing some suppression (Supplemental 6.1, Panels B and C).

Next, we examined whether this suppression was seen in the full-length protein containing a single mutation in the PH domain, or in expression of just the PH domain with the single mutation (Supplemental 6.1, Panels D-F). *CDC42* over-expression was able to suppress the temperature sensitivity of both the *boi1PH-W794R* and the *boi1PH-F799S* mutants, and only *RHO3* over-expression showed some suppression of the *boi1PH-W794R*.

We were surprised to find that the original conditional *boi1* mutants show temperature-sensitive growth at 35°C, yet grew quite well at 37°C (see Figure 5B). It has been shown that shifting yeast cells from 26°C to 37°C elevates the level of the plasma membrane signaling lipid, PI(4,5)P_2_ (Stefan et al., 2002). Moreover, the mutations in the PH domain that confer temperature sensitivity likely affect the binding of PI(4,5)P_2_ (See Figure 5F). If this is the reason why the conditional mutants grew better at 37°C, we predicted that over-expression of the enzyme that synthesizes PI(4,5)P_2_ should suppress growth at 35°C. Accordingly, we over-expressed the PI-4P 5-kinase *MSS4* in each of the three conditional mutants, and found that all were partially suppressed, but to different degrees (Supplement 6.2). Next, we examined cells expressing full length Boi1 with either the single W794R or F799S mutations, or these mutations in the context of the Boi1 PH domain (Supplemental 6.2). In both cases, *MSS4* over-expression suppressed the constructs harboring the W794R mutation, but not those containing the more severe F799S mutation (Supplemental 6.2). Interestingly, at 37°C, *MSS4* over-expression suppressed the mutations in the context of the full-length protein, but not in the context of the PH domain. This finding is addressed later in the Discussion.

### Boi1 and Bem1 have a secondary overlapping essential function that requires Boi1’s SH3 domain

The redundant Boi1 and Boi2 proteins were initially identified in a screen for interactors of Bem1’s second SH3 domain (SH3b) (Bender et al., 1996; Matsui et al., 1996). Disrupting the function of Boi1’s SH3 domain with the *boi1W53K* mutation in a *boi2*Δ background rendered *BEM1* essential. Therefore, in *boi2*Δ cells there is an essential function requiring either Boi1-SH3 or Bem1 (Matsui et al., 1996). We confirmed that the *boi1W53K* mutation does not confer a growth defect in an otherwise wildtype background, or in *boi2*Δ cells, and that *boi2*Δ *boi1-W53K bem1*Δ is lethal (Figure 7A, B). We also confirmed that *bem1*Δ and *bem1*Δ *boi2*Δ are temperature sensitive (Figure 7A).

**Figure 7:**
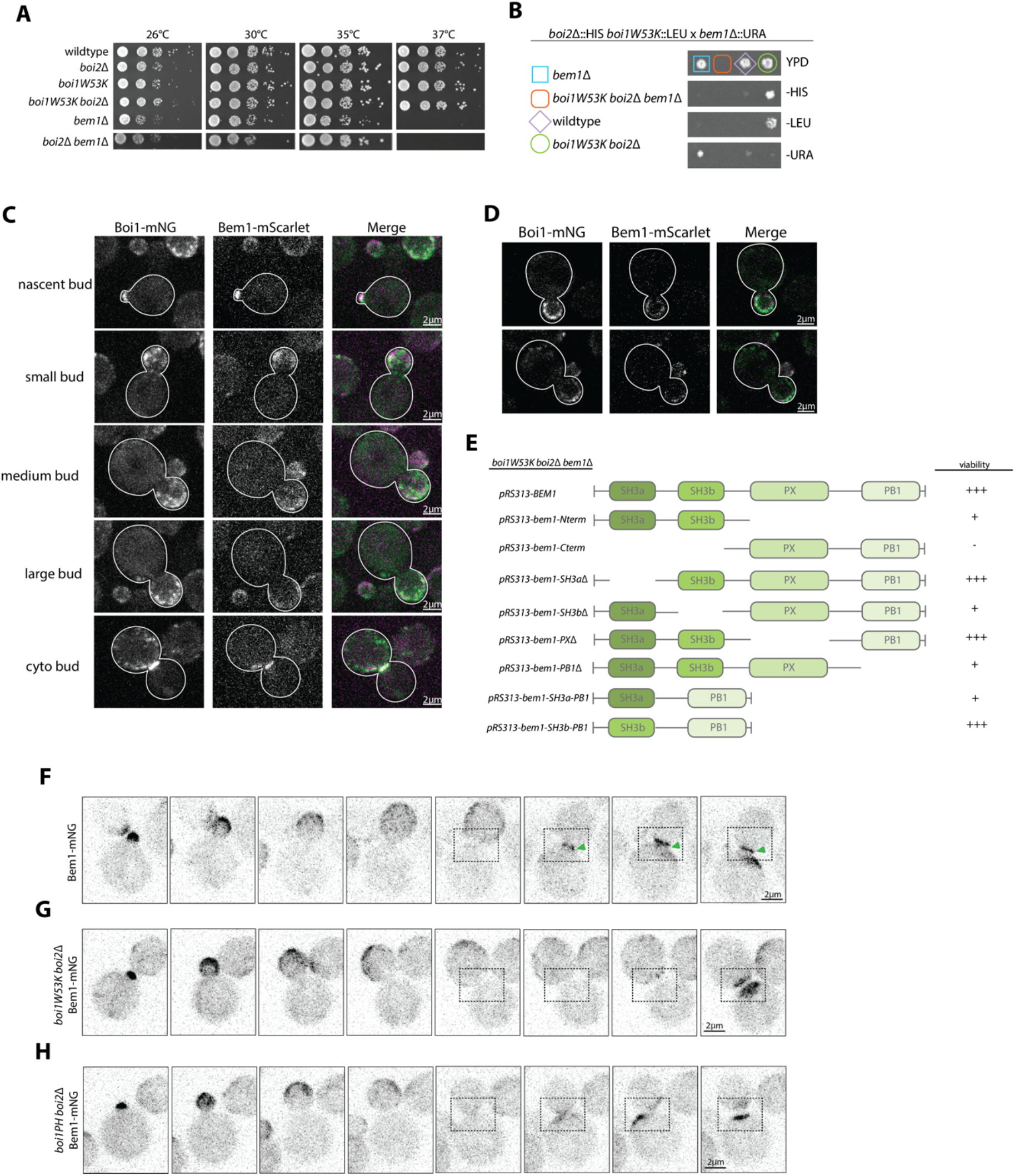
Bem1 and Boi1 have a secondary essential overlapping function that requires Boi1’s SH3 domain. (A) Growth assays of wildtype, *boi2Δ, boi1W53KΔ, boi1W53K boi2Δ, bem1Δ*, and *boi2Δ bem1Δ*. Cells were grown on YPD plates for two days at the respective temperatures. (B) Tetrad dissections of *boi2Δ boi1W53K* crossed with *boi2Δ bem1Δ*. Dissections revealed that *boi1W53K boi2Δ bem1Δ* is inviable. Spores are representative of 3 plates of dissections. Spores were grown on respective media for 2 days. (C) Microscopy images of Boi1-mNG and Bem1-mScarlet at five different stages of bud growth including cytokinetic bud. Microscopy was max projections of 15 planes covering ∼4.2µm of the cell. 300ms exposures were used to visualize Boi1-mNG and Bem1-mScarlet. Scale bars are 2µm. (D) Individual slices from two of the panels from C showing a single plane in the middle of the cell. Images were taken the same as in C, and scale bars are 2µm. (E) Domain map of Bem1 truncations and their growth in *boi1W53K boi2Δ* cells. Growth is indicated as follows: none (-), temperature sensitive (+), or wildtype-like (+++). Growth assays shown in Supplemental Figure 6.2 (F, G, and H) Images of: Bem1-mNG, *boi1W53K boi2Δ* Bem1-mNG, and *Boi1PH boi2Δ* Bem1-mNG, respectively. Images are max projections of 15 z-planes throughout the cell ∼4.2µm. Timelapses were taken with 5 minute intervals, with 100ms exposure. All scale bars are 2µm.

While it is known that Boi1/2 and Bem1 each localize to sites of polarization, it is not clear how well they co-localize. We therefore imaged Boi1-mNG and Bem1-mScarlet and found that they colocalize at every stage in the cell cycle (Figure 7C and D). However, Boi1 arrives at the bud neck about a minute before Bem1 when compared to the cytokinesis marker Myo1 (compare Figure 1E to Supplemental Figure 7.1). We also observed Bem1 arriving after Boi1 in dual-color microscopy videos (Video 7). Furthermore, in *boi1W53K boi2*Δ or *boi1PH boi2*Δ cells, Bem1-mNG is not recruited to the bud neck (Figure 7F-H). These results indicate that Bem1 is recruited to the bud neck by Boi1 through its SH3 domain. Since *boi1W53K boi2*Δ has no growth or cytokinesis defect, this implies that Bem1 does not play an important role in cytokinesis.

To define the minimal Bem1 domains necessary for growth in a *boi1W53K boi2*Δ background, we conducted a plasmid shuffle screen with several truncated versions of Bem1. A construct lacking both SH3 domains was not viable, and lacking just the SH3a grew well, but lacking SH3b grew poorly (Figure 8E, Supplemental 7.2). Further analysis showed that in the *boi1W53K boi2*Δ background, robust growth required the presence of both the SH3b and PB1 domains. Indeed, a fusion of SH3b-PB1 restored growth to the wildtype levels, whereas a fusion of SH3a-PB1 did not. Moreover, the SH3b-PB1 construct fully suppressed the temperature sensitivity at 37°C of *bem1*Δ (Figure 7E, Supplemental 7.2).

**Figure 8:**
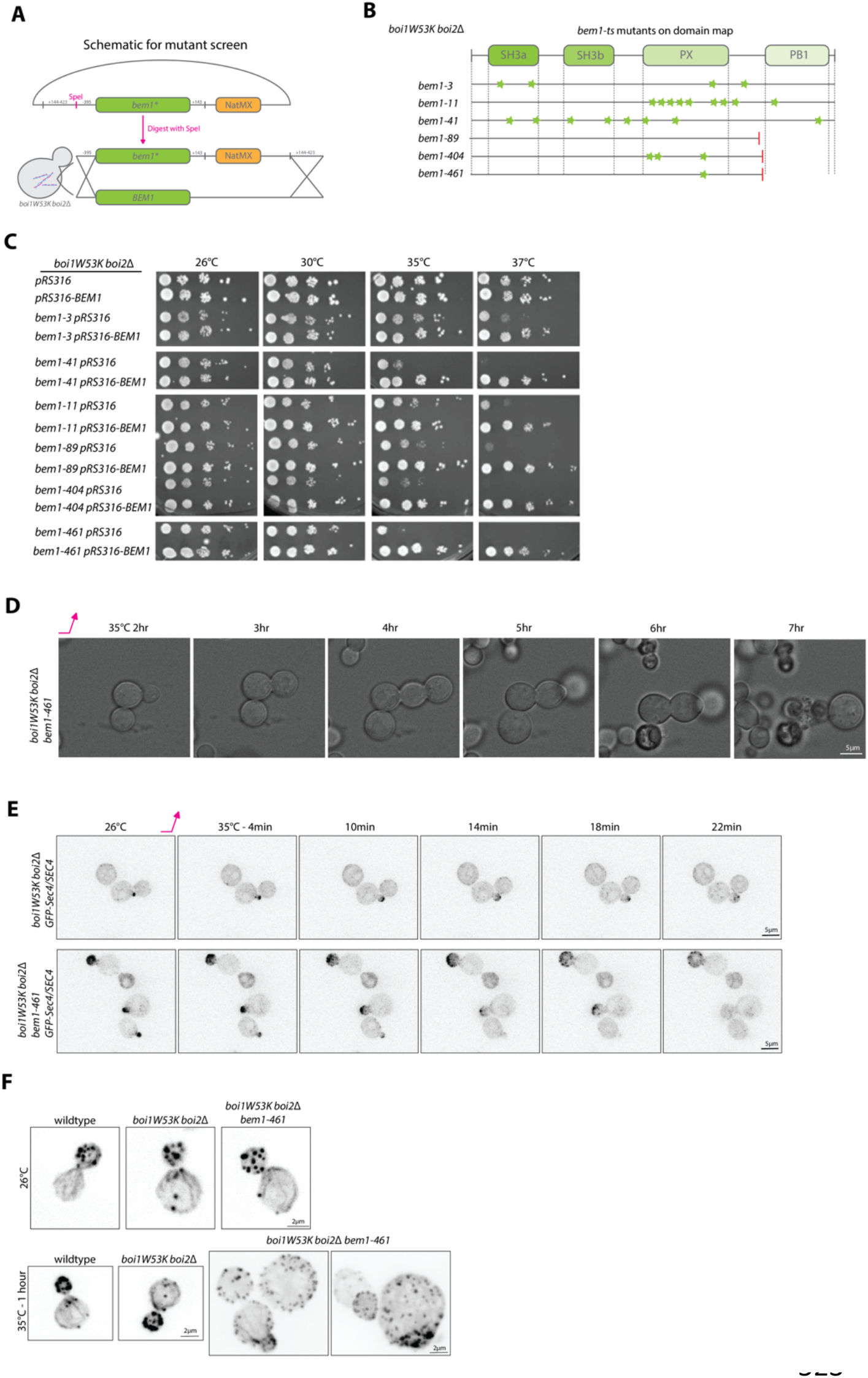
*BEM1* conditional mutants exhibit unusual cellular morphology, budding pattern, and actin morphology. (A) Schematic for the integration of mutated *BEM1* to replace the endogenous *BEM1*. (B) Domain map of Bem1 showing where several of the mutations from different candidates fall. (C) Growth assay of bem1 conditional mutants with empty pRS316 or a pRS316-BEM1 plasmid grown at 26°, 30°, 35°, or 37°C for 3 days. (D) DIC images of *bem1-461 boi1W53K boi2Δ* at 26°C and 37°C over the course of a timelapse up to 7 hours of growth. Scale bars are 5µm. (E) Upper panel is *boi1W53K boi2Δ GFP-Sec4/SEC4* diploid cells at 26°C and then shifted to 35°C for ∼22 minutes. Lower panel shows the *bem1-461 boi1W53K boi2Δ GFP-SEC4/SEC4* diploid strain. Scale bars are 5µm. (F) Actin phalloidin staining of the cells grown at 26°C or after shifting to 35°C for 1 hour. Images are max projections of 15 z-planes with 100ms exposure. Scale bars are 2µm.

We conclude that the SH3b and PB1 domains of Bem1 can fully substitute for the full-length protein, and that the SH3b domain performs a redundant function with the Boi1 SH3 domain. Additionally, in the *Boi1PH boi2*Δ *bem1*Δ strain, the Bem1-SH3b-PB1 construct could restore growth comparable to full length *BEM1* (Supplemental 7.2). Thus, the three genes *BOI1, BOI2*, and *BEM1* can be functionally replaced by the *Boi1PH* and Bem1-SH3b-PB1 constructs.

### Conditional *bem1* mutants generated in the *boi1W53K boi2*Δ background have significant morphological and budding defects

To facilitate investigation of the essential function of Bem1 in *boi1W53K boi2*Δ cells, we targeted mutated *BEM1* to its chromosomal locus and screened for temperature sensitive mutations (Figure 8A). Several candidates were recovered and all were found to be recessive to *BEM1* (Figure 8B and C). Sequence analysis showed many mutations in some of the conditional *bem1* alleles, but notably several of them were stop codons that eliminated the important PB1 domain (Supplemental 8.1). Two of the mutants that had truncations in the PB1 domain also had mutations in the PX domain which caused a more severe temperature sensitivity compared to the truncation alone (Figure 8C, Supplemental 7.2).

One of these mutants, *bem1-461* was selected for further study. Upon shifting to 37°C, the mutant continued to grow but in a largely isotropic manner, and then lysed. The lysing pattern is similar to what we saw in the conditional *boi1* mutants; however, growth is not initially impaired. Additional other phenotypes result including bud-on-bud pattern similar to mutants of *pkc1-2* (Paravicini et al., 1992)(Figure 8C, Supplemental 8.2). This mutant also showed a largely depolarized actin cytoskeletal structure accounting for the severe defects in polarized growth (Figure 8E). Replacement of *SEC4* with *GFP-SEC4* suppressed the temperature sensitivity of the *bem1-ts*, so we examined its distribution in *SEC4/GFP-SEC4* diploids (Figure 8D). The secretory vesicles marked by *GFP-SEC4* do not initially depolarize upon shifting to the restrictive temperature, but eventually do. This is consistent with the depolarized actin cytoskeleton and depolarized growth seen after shifting for an extended period. Thus, the onset of defects appears to be quite slow in this mutant, making pinpointing a specific process difficult.

## Discussion

Boi1/2 have been implicated in functioning during cell polarity by interacting with Cdc42, PIP_2_, the exocyst and Sec1 during exocytosis, and actin nucleators. Genetic analysis has also clearly shown that Boi1/2 share an overlapping and essential function with Bem1. With this broad set of proposed roles, we aimed to probe the essential functions of Boi1/2 and Bem1 by analyzing each protein’s localization throughout the cell cycle, and by generating and characterizing conditional mutants in the essential domains.

Careful characterization showed that both Boi1 and Boi2 are present in dynamic cortical patches where Boi2 patches are more stable. Surprisingly, although Boi1 and Boi2 are functionally redundant, they frequently showed distinct patches in the bud, suggesting that they may have divergent as well as overlapping functions. It is unclear whether Boi1 or Boi2 may be arriving in a sequential order, as examples of the two proteins arriving one before the other have been observed, and it is unclear whether it is to the same site on the membrane, or rather nearby. Both proteins localize at the bud neck prior to cytokinesis. This highlights a role at the neck to potentially aid in the organization of cytokinetic proteins or to allow for exocytic vesicles to accumulate at the neck to ensure the closure of the plasma membrane and cell wall during separation.

Given the potential interplay of Boi1/2 in actin structure and secretory vesicle exocytosis, we examined the effect of disrupting both actin and exocytosis on Boi1/2 localization. Neither had any affect, showing that if Boi1/2 play roles in these processes, they are upstream in the pathway. Our analysis found that Boi1/2 remains at the cell cortex upon shifting *sec6-4* to the restrictive temperature, again suggesting that they may function upstream of exocytosis. We also asked if Boi1/2 patches correlate with sites of secretory vesicle exocytosis especially given that Boi1/2 patches have a lifetime about double that of secretory vesicle tethering (Gingras et al., 2022). Interestingly, there was very little colocalization between Boi1/2 and Sec4, and since Boi2 patches do not move extensively on the membrane, it is difficult to reconcile with a model that invokes a direct and essential role for Boi1/2 in the tethering/exocytosis of secretory vesicles. Moreover, secretory mutants show immediate cessation of growth upon shifting to the restrictive temperature, which was not seen in our boi1 conditional mutants, again shedding doubt that they play an essential role in exocytosis. Nevertheless, two studies have made a clear case for a role in exocytosis (Kustermann et al., 2017; Masgrau et al., 2017). In both studies the level of Boi1 proteins was depleted by either dilution or targeted degradation, both of which take significant time. It is possible that secretory vesicle accumulation in both these studies is a secondary and indirect effect of inhibiting another important process. As the essential region of Boi1/2 lies in the C-terminal PH domain, we turned to examining the properties of cells that express just the Boi1-PH domain in *boi2*Δ cells.

The *Boi1-PH boi2*Δ cells grew normally at all temperatures, and localized tightly to the cortex of the bud. The localization was more diffuse than Boi1-mNG, indicating that the part of the protein lacking in this construct is responsible for patch localization. Boi1-PH-mNG did not relocate to the bud neck for cytokinesis, whereas the non-essential regions containing the SH3 domain did, indicating that the role of Boi1/2 in cytokinesis is not important for cell viability.

It is clear from these studies that the essential function of the Boi1/2 proteins is not simple, and therefore we set out to generate conditional mutations in *BOI1* in a *boi2*Δ strain. Generating conditional mutations in *BOI1* turned out to be challenging, and the construction of the *boi1-ts* library and mutants were the first published attempts. These conditional mutations were only viable on a plasmid, presumably due to a slightly higher level of expression than when placed in the chromosome. These mutants had an interesting phenotype in that they grew at 26°C, but not at 35°C, but did again at 37°C. Moreover, due to the delicate nature of the mutants, they would not tolerate tagging of key proteins involved in secretion, such as Sec4 and subunits of the exocyst. This condition made it difficult to characterize the mutants to understand which cellular process the *boi1-ts* were involved in. Furthermore, the fact that the conditional mutants continued to exhibit cell growth at the restrictive temperature, unlike exocytic mutants *sec6-4* or *sec4-8*, indicates that the essential function is unlikely to be directly involved in exocytosis. Thus, the characterization did not give a clear phenotype as we had hoped – having apparent defects in growth and eventually lysing after extended periods at the restrictive temperature.

In previous unpublished work, we isolated conditional mutations in *BOI2* in a *boi1*Δ background. Many of the *BOI1* and *BOI2* conditional mutants had mutations in either of two homologous residues: W794R or F799S in *BOI1*, or W786R or 791S in *BOI2*. Replacing the *BOI1* gene in a *boi2*Δ strain with the single W794R or F799S mutation into full length Boi1, or Boi1-PH, resulted in temperature-sensitive growth. Moreover, a screen in which any amino acid was placed at position 794 or 799 revealed the same amino acid changes already recovered that conferred temperature sensitivity. Thus, our mutational analysis reveals the critical nature of these two residues, which structurally are localized near the predicted PIP_2_ binding region of the PH domain. Consistent with this, over-expression of the kinase that generates PIP_2_ at the plasma membrane, Mss4, partially suppresses the conditional mutants at 35°C.

As noted above, our temperature sensitive *boi1* mutants, failed to grow at 35°C, but did grow at 37°C. We suspected that this phenotype might be due to a defect in PIP_2_ binding, as PIP_2_ levels rise upon shifting to 37°C (Stefan et al., 2002). Interestingly, the *boi1-PHW794R* allele in a *boi2Δ* background is temperature sensitive for growth at both 35°C and 37°C. While *MSS4* over-expression partially restores growth at 35°C, it does not do so at 37°C. This suggests that while elevating PIP_2_ may help growth in the presence of *boi1-PHW794R*, it is not the only essential function of the PH domain. Since the same mutation in the context of the full length Boi1 protein does grow at 37°C, this essential function of the PH domain at 37°C must be at least partially redundant as it can be satisfied by some other part of the full-length molecule.

As far as we are aware, this is the only incidence where a single PH domain has been shown to be essential. Simply binding a lipid cannot be essential itself, so it is necessary to predict that the Boi1PH construct must bind another factor, perhaps acting as a coincidence detector. An obvious candidate is Cdc42 that has been shown to bind the PH domain (Bender et al., 1996) and *CDC42* over-expression, or its activator *CDC24*, can suppress our conditional mutants. However, arguing against this model is the generation of the triple R827A, L829A, T894A mutant in the Boi1PH domain that no longer binds Cdc42 and that has no effect on cell growth (Kustermann et al., 2017). Thus, the essential function of the PH domain remains open, and perhaps genetic approaches using the new conditional mutants will be more illuminating.

We next turned our attention to the overlapping function between Boi1/2 and Bem1. We confirmed that Bem1 becomes essential in a *boi1W53K boi2Δ* strain. By deletion analysis, we found that fusing the second SH3 domain of Bem1 to its C-terminal PB1 domain could provide growth equivalent to wildtype cells. Thus, the two essential functions of Bio1/2/Bem1 can be supplied by just the Bem1-SH3-PB1 construct replacing *BEM1*, and the Boi1PH replacing *BOI1* and *BOI2*.

To analyze the essential function of Bem1 in the *boi1W53K boi2Δ* cells, we once again turned to generating conditional mutations in *BEM1*. In contrast to the case described above, generating conditional *BEM1* mutations by direct insertion into the chromosome was relatively straightforward. Upon shifting to the restrictive temperature, one of these mutants, *bem1-461*, continued to grow isotropically and then lysed. Consistent with depolarized growth, the actin cytoskeleton was highly depolarized after shifting to the restrictive temperature for an hour. However, the onset of these phenotypes was slow - *GFP-Sec4* marked secretory vesicles did not depolarize immediately upon shifting to the restrictive temperature, so is difficult to assess if the depolarized phenotype is a direct or indirect consequence of affecting the essential function of Bem1.

In this study, we have defined more clearly the localization and dynamics of the Boi1/2 scaffolding proteins and identified two critical residues that impact the essential function of the Boi1/2 PH domains. We have also clarified the regions, the SH3 and PB1 domain, important in Bem1 that provide full function in a *boi1W53K boi2Δ* cell. It is somewhat disappointing that the phenotypes of the two sets of conditional mutations that we generated in *BOI1* and *BEM1* where not more illuminating, but future genetic approaches using these mutants may help in pinpointing the precise essential functions of these scaffolding proteins.

## Data Availability

All strains and plasmids are available upon request. The authors affirm that all data important and necessary for the conclusions within this article are present within the article, figures, and Tables 1 and 2.

## Acknowledgements

The authors would like to thank members of the Bretscher lab for helpful comments and suggestions pertaining to this work: Dr. MJ Shin, Dr. Andrew Lombardo, Riasat Zaman, David McDermitt, and Valerie Awad. Thank you to Dr. Alan Sulpizio from the Mao lab at Cornell, and Dr. Chris Fromme, and Dr. Thomas Fox for helpful comments and suggestions, and to Dr. Scott Emr and Dr. Jeffrey Jorgensen for supplying the *MSS4*-containing plasmids used in this study. This study was supported by NIH grants RO1GM39066 and R35GM131751. Authors declare no conflict of interest.

A. Sulpizio conducted experiments, wrote this research, and conceptualize the project. L. Herpin conducted experiments pertaining to the Bem1 research and helped conceptualize the project. Dr. Gingras helped with truncations of Boi1, and with microscopy. Dr. Liu conducted the Boi2 conditional mutant screen, and Dr. Bretscher conceptualized the project and wrote this research.

## Materials and Methods

### Development of Yeast Strains and Growth in Different Media Types

Yeast strains used in this study are listed in ***Table 1***. All strains used were haploid BY4741/2 unless otherwise noted. Yeast were grown in either selectable synthetic complete liquid media or on plates. Plates supplemented with 5-Fluoroorotic Acid (5FOA) were used to shuffle out yeast containing a plasmid with URA (Boeke et al., 1984). Sporulation of yeast was conducted by growing strains at 26°C in sporulation media (1% yeast extract, 1% potassium acetate, and 0.05% glucose). Tetrad dissections were performed using a Zeiss Standard 14 dissection scope.

### Molecular Cloning

Deletion of genes or tagging was performed using PCR-constructed DNA fragments with 30bps of homology to the chromosome (Longtine et al., 1998). Truncations of Boi1 N-terminal region was also made through homologous recombination of mNG fragment to the end of each domain. Plasmids used in this study are listed in ***Table 2***. Truncated Boi1PH(730-980) was made by cloning Boi1PH into integrating vector: pRS305 with Boi1 downstream region (+150-350bp) cloned ahead of the Boi1 promoter (-370bp) with a SalI endonuclease cut site separating the two fragments. Upon digestion with SalI, the linearized vector completely replaces the endogenous gene with LEU selection (Figure 5A) (Chernyakov et al., 2013). Other integration plasmids were used for tagging proteins for fluorescent microscopy: pRS30Nat-mScarlet-Sec4 adapted from (Donovan & Bretscher, 2012), pRS306-GFP-Sec4 (same study), pRS306-Bni1-GFP (Pruyne). Point mutants on plasmids were made by developing back-to-back 5’ phosphorylated primers with one containing the mutation in the middle of the sequence. Protocol was adapted from Thermo Scientific Phusion Site-Directed Mutagenesis Kit. The point mutant: *boi1W53K* was made by integration of *pRS305-boi1W53K* which has a 300bp homologous region to upstream of Boi1’s promoter (685-385bp upstream of start codon) cloned after the 3’UTR with a XbaI endonuclease cut site. Upon linearization of the plasmid, the *boi1W53K* mutant replaced the endogenous gene. Yeast transformations were conducted using a standard Lithium Acetate protocol. Bacterial cloning techniques used either Top10 or DH5alpha competent cells.

### Construction of Temperature Sensitive Mutants

Temperature sensitive mutants were made through error-prone PCR mutagenesis as adapted (Cadwell & Joyce, 1992; Wilson & Keefe, 2001). The mutated fragment of *BOI1* was cloned into either a pRS315 or pRS305 vector for further experiments. At least 3 separate libraries were constructed for each strategy collecting bacterial clones off of agar plates and Maxi prepping the libraries. Using Strategy 1, (and also described above) the *boi1-ts* fragment was transformed into cells (Figure 5A). Using Strategy 2, *boi1-ts* was transformed into cell types with *boi1*Δ *boi2*Δ and *pRS316-BOI1*. The fragment of *boi1-ts* had differing homologous regions to the chromosome and not to the *BOI1* fragment on the pRS316 plasmid so as not to recombine with the plasmid. Transformations were plated on -LEU plates (Strategy 1) or -LEU/-URA plates (Strategy 2) and then placed at 26°C for 2 days. For Strategy 1: Plates were replicated using velvet squares and replicated plates were placed at 35°C for 2 days, while the original plate was placed back at 26°C. The two plates were then screened by eye to find colonies capable of growth at 26°C and that were dead at 35°C. Over 15,000 colonies were screened and candidates were selected for further dilution assays on selectable media. For Strategy 2: Plates were replicated onto -LEU 5FOA media to eliminate the *pRS316-BOI1* plasmid and were grown for 3 days at 26°C. After sufficient growth, they were then replicated again onto two -LEU plates, one placed at 26°C and the other at 35°C for 2 days. They were then screened by eye to find temperature sensitive colonies. *Bem1-ts* libraries were made using the same strategy (Figure 7F).

*BOI2* conditional mutants were made through PCR-mutagenized DNA encoding full length *BOI2*. The mutagenic PCR was performed using the following mixture: 1mM dCTP, 1mM DTTP, 0.2mM dATP, 0.2mM dGTP, 75ng template DNA (pWL106), 5.5mM MgCl_2_, 0.5mM primers, 1X Taq buffer (Roche), 10µg BSA, and 0.05% Tween-20 per 100µL reaction. The mutagenic PCR was subjected to eight cycles of amplification and the BamHI-NotI fragment was subcloned into pWL105 to make the libraries. Transformants were grown at the permissive temperature of 26°C and then replica-plated to 37°C to identify temperature sensitive clones. 14,000 colonies were screen and 5 temperature sensitive strains were saved (Figure 5, Supplemental 4.1, and Table 1).

### Microscopy Techniques and Imaging

For live-cell imaging, yeast were grown in liquid culture to mid-log phase (0.5-0.8OD) and were prepared by adhering 200µL of cells to 4-well glass-bottom dishes (CellVis) coated with concanavalin-A submerged with proper media for cell survival. For fixed imaging, yeast were grown in liquid culture to mid-log phase and 4% paraformaldehyde was added to the culture for 10 minutes. Cells then followed appropriate protocol (See Actin Staining) and then were adhered to coverslips, again by using concanavalin-A. Coverslips were then mounted using soft mounting media (Invitrogen) and secured by using clear nail polish. Some fixed cells were also imaged using agarose pads, where 4µL of fixed cells were added to a 1% agarose pad and media was allowed to diffuse into the agarose (Pringle et al., 1989; Shin et al., 2018). All fluorescent microscopy was conducted on an inverted microscope (Leica DMI6000B) which had a spinning disk confocal unit (Yokogawa CSU-X1) with 100 × 1.45 NA objective (Leica) with either a Evolve 512Delta EMCCD or with a Flash 4.0v2 CMOS camera. DIC imaging was conducted on this microscope as well as on a Leica De-Convolution Microscope (DMi8). Slidebook 6.0 or FIJI software were used for image analysis.

### Cell Death Assay

Yeast cells were analyzed for death rate in Supplemental 5.1 during live-cell imaging. Cells were marked “dead” when lysing took place, and was measured to verify death using Live-or-Dye (Biotium). Dye was in 10% DMSO and was added 1:100 into yeast cells and allowed to incubate for at least 30 minutes before shifting the temperature. After confirming that the yeast lysing was indeed killing the cells, we used DIC to measure the rest of the cultures.

### Fluorescence Recovery after Photobleaching (FRAP)

For the FRAP experiment, specific regions of the cell, either a Boi1/2 patch or half of the bud neck, were photobleached using a single point with the 3i Vector photobleaching system after 1s of capturing frames were taken every 160ms for 32s. Photobleaching control was taken as the region of a neighboring mother cell. In analysis graphs background control was subtracted and normalized to max intensity.

### CherryTemp Mutant Microscopy

Imaging the temperature sensitive mutants using traditional methods is highly variable and difficult to control. In these studies, we used a microfluidic device temperature control system fitted to either the 3i Confocal or the Leica microscopes (CherryBiotech). Cells were adhered to a glass slide using concanavalin A, a plastic spacer with a cut-out hole was added on top of the cells, and 50µL of selectable media was placed on top of the cells. The ThermoChip Immersion was placed on top of the cells with the plastic spacer so the yeast were close to the microfluidic chamber and still had space to continue growth. The CherryTemp was set to 26°C and shifted to 35°C depending on the experiment. The temperature shift takes only seconds, but we waited 5min for equilibration and then started imaging.

### Actin Staining

The actin staining was conducted using two different protocols (noted in Figure legends). The first staining protocol washed the fixed cells with PBS x2, and then with PBS+0.1%TritonX. Cells were spun down after each wash on a very low centrifugation speed – 500g. The cells were incubated with PBS+0.1%TritonX for 20min, and then spun down and were incubated with 4% phalloidin AlexaFlour-488 in PBS+0.1%TritonX. Prior to incubation, the phalloidin was dried and resuspended in equal PBS+0.1%TritonX. After phalloidin incubation cells were spun down and washed with PBS+0.1%TritonX x2, and once with PBS. 1:1000 DAPI incubation was also conducted for 10 minutes prior to adhering to the concanavalin-a cover slips and mounted using soft mounting pro-diamond gold (Invitrogen) onto glass slides (Figure 3A). The second phalloidin stain protocol used dried phalloidin AlexaFlour-568 resuspended in PBS+0.1%TritonX. 4µL of fixed cells were added 3x to 1% agarose pads until the media diffused into the agarose. Phalloidin was added on top of the cells and sufficient time was allotted for the PBS to diffuse (2-3 minutes). Once diffused, cover-slips were added on top of the samples and then they were imaged.

### Latrunculin-A and CK-666 Treatment

Treatment of the cells with Latrunculin A (LatA) was conducted prior to fixation for actin staining. Cells were treated to a final concentration of LatA of 250µM in 2% DMSO (Thermo Fisher) for 5min before addition of 4% PFA. For live cell imaging with LatA cells were adhered to the glass-bottom dishes and then LatA was spiked into the media with a final concentration of 250µM in 2% DMSO. Control cells were treated with just 2% DMSO (Ayscough et al., 1997). For the CK-666 treatment the cells were incubated with CK-666 for 5 minutes prior to fixation or prior to adhering for live cell imaging.

## Notes

### Competing Interest Statement

The authors have declared no competing interest.

