## Supplemental Material for "Generation and characterization of conditional yeast mutants affecting each of the two essential functions of the scaffolding proteins Boi1/2 and Bem1"

**Table 1: List of Yeast Strains**

| ***Strain Genotype*** | ***Name*** | ***Reference*** |
|---|---|---|
| *Mat-a BY4741 Wildtype* | ABY1655 | C. Boone |
| *Mat-$\alpha$ BY4742 Wildtype* | ABY1656 | C. Boone |
| *Mat-a boi1Δ::URA3* | ABY4643 | This study |
| *Mat-$\alpha$ boi1Δ::URA3* | ABY4644 | This study |
| *Mat-a boi2Δ::HISMX* | ABY4645 | This study |
| *Mat-$\alpha$ boi2Δ::HISMX* | ABY4646 | This study |
| *Mat-a Boi1-mNG::URA3* | ABY4647 | This study |
| *Mat-a Boi2-mNG::HISMX* | ABY4642 | This study |
| *Mat-a Boi1-mNG::URA3 Myo1-mScarlet::NATMX* | ABY5021 | This study |
| *Mat-a Boi1-mNG::URA3 Boi2-mScarlet::NATMX* | ABY5031 | This study |
| *Mat-a tpm1-2::LEU2 tpm2Δ::HISMX* | ABY944 | D. Pruyne |
| *Mat-$\alpha$ tpm1-2::LEU2 tpm2Δ::KANMX Boi2-mNG::HISMX* | ABY5029 | This study |
| *Mat-a Boi2-mNG::HISMX mScarlet-Sec4::NATMX* | ABY5028 | This study |
| *Mat-a boi1Δ::URA3 Boi2-mNG::HISMX mScarlet-Sec4::NATMX* | ABY5040 | This study |
| *Mat-a sec4-8::HISMX* | ABY9429 | P. Novick |
| *Mat-a sec4-8::HISMX Boi1-mNG::URA3* | ABY5034 | This study |
| *Mat-a sec4-8::HISMX Boi2-mNG::HISMX* | ABY5036 | This study |
| *Mat-a sec6-4::HISMX* | ABY9035 | P. Novick |
| *Mat-a sec6-4::HISMX Boi1-mNG::URA3* | ABY5035 | This study |
| *Mat-a sec6-4::HISMX Boi2-mNG::HISMX* | ABY5037 | This study |
| *Mat-$\alpha$ Boi1PH::LEU2 boi2Δ::HISMX* | ABY4770 | This study |
| *Mat-$\alpha$ Boi1PH-mNG::NATMX boi2Δ::HISMX* | ABY4772 | This study |
| *Mat-a Boi1SH3-mNG::LEU2* | ABY4774 | This study |
| *Mat-a Boi1SH3-SAM-mNG::LEU2* | ABY4775 | This study |
| *Mat-a Boi1SH3-SAM-PxxP-mNG::LEU2* | ABY4776 | This study |
| *Mat-a boi1Δ::HISMX boi2Δ::KANMX pRS315-boi1-1* | ABY4734 | This study |
| *Mat-$\alpha$ boi1Δ::HISMX boi2Δ::KANMX pRS315-boi1-2* | ABY4735 | This study |

| | | |
|---|---|---|
| *Mat-α boi1Δ::HISMX boi2Δ::KANMX pRS315-boi1-3* | ABY4702 | This study |
| *Mat-a boi1Δ::HISMX boi2Δ::KANMX pRS315-BOI1* | ABY4737 | This study |
| *Mat-a boi1Δ::HISMX boi2Δ::KANMX pRS315-Boi1PH* | ABY4738 | This study |
| *Mat-a boi1Δ::KANMX boi2-1::LEU2* | ABY3208 | W. Liu |
| *Mat-a boi1Δ::KANMX boi2-4::LEU2* | ABY3211 | W. Liu |
| *Mat-a boi1Δ::KANMX boi2-5::LEU2* | ABY3212 | W. Liu |
| *Mat-a boi1Δ::KANMX boi2-6::LEU2* | ABY3213 | W. Liu |
| *Mat-a boi1Δ::KANMX boi2-7::LEU2* | ABY3214 | W. Liu |
| *Mat-a boi1Δ::HISMX boi2Δ::KANMX pRS315-boi1W794R* | ABY4766 | This study |
| *Mat-a boi1Δ::HISMX boi2Δ::KANMX pRS315-boi1F799S* | ABY4764 | This study |
| *Mat-a boi1Δ::HISMX boi2Δ::KANMX pRS315-boi1PHW794R* | ABY4905 | This study |
| *Mat-a boi1Δ::HISMX boi2Δ::KANMX pRS315-boi1PHF799S* | ABY4909 | This study |
| *pck1Δ::LEU2 yEP50-pck1-2 SEY6210* | AAY603 | S. Emr |
| *Mat-a boi1W53K::LEU2 boi2Δ::KANMX* | ABY4110 | This study |
| *Mat-a bem1Δ::URA3* | ABY4102 | This study |
| *Mat-α bem1Δ::URA3 boi2Δ::HIS* | ABY4107 | This study |
| *Mat-a boi1-mNG::URA3 Bem1-mScarlet::NATMX* | ABY4942 | This study |
| *Mat-a bem1-mNG::URA3* | ABY4140 | This study |
| *Mat-a bem1-mNG::URA3 Myo1-mScarlet::NATMX* | ABY4175 | This study |
| *Mat-α boi1W53K::LEU2 boi2Δ::HISMX Bem1-mNG::URA* | ABY4139 | This study |
| *Mat-α Boi1PH::LEU2 boi2Δ::HISMX Bem1-mNG::URA* | ABY4148 | This study |
| *Mat-a boi1W53K::LEU2 boi2Δ::KANMX bem1Δ::NATMX pRS313-BEM1* | ABY4114 | This study |
| *Mat-α boi1W53K::LEU2 boi2Δ::HIS bem1-3::NATMX* | ABY4116 | This study |
| *Mat-α boi1W53K::LEU2 boi2Δ::HISMX bem1-11::NATMX* | ABY4119 | This study |
| *Mat-α boi1W53K::LEU2 boi2Δ::HISMX bem1-41::NATMX* | ABY4122 | This study |
| *Mat-α boi1W53K::LEU2 boi2Δ::HISMX bem1-89::NATMX* | ABY4128 | This study |
| *Mat-α boi1W53K::LEU2 boi2Δ::HISMX bem1-404::NATMX* | ABY4134 | This study |
| *Mat-α boi1W53K::LEU2 boi2Δ::HISMX bem1-461::NATMX* | ABY4136 | This study |

| | | |
|---|---|---|
| *Mat-a/$\alpha$ boi1W53K::LEU2 boi2Δ::HISMX bem1-461::NATMX GFP-Sec4::URA3/SEC4* | ABY4904 | This study |

**Table 2: List of plasmids**

| *Plasmid Name* | *Bacterial Strain* | *Name* | *Reference* |
|---|---|---|---|
| *pRS315-BOI1* | Top10 | AB4538 | This study |
| *pRS316-BOI1* | Top10 | AB4452 | This study |
| *pRS316-BOI2* | Top10 | AB4446 | This study |
| *pRS315-boi1-1* | Top10 | AB4590 | This study |
| *pRS315-boi1-2* | Top10 | AB4591 | This study |
| *pRS315-boi1-3* | DH5$\alpha$ | AB4565 | This study |
| *pRS315-boi1W794R/F799S* | DH5$\alpha$ | AB4729 | This study |
| *pRS315-boi1PH* | Top10 | AB4576 | This study |
| *pRS315-boi1PHW794R* | DH5$\alpha$ | AB4730 | This study |
| *pRS315-boi1PHF799S* | DH5$\alpha$ | AB4732 | This study |
| *pRS315-boi1PHW794R/F799S* | DH5$\alpha$ | AB4731 | This study |
| *yEP352-Cdc42* | DH5$\alpha$ | AB3929 | R. Gingras |
| *yEP352-Rho1* | DH5$\alpha$ | AB3920 | R. Gingras |
| *yEP352-Rho2* | DH5$\alpha$ | AB3922 | R. Gingras |
| *yEP352-Rho3* | DH5$\alpha$ | AB3865 | R. Gingras |
| *yEP352-Rho4* | DH5$\alpha$ | AB3925 | R. Gingras |
| *yEP352-Rho5* | DH5$\alpha$ | AB3927 | R. Gingras |
| *pRS426-MSS4* | DH5$\alpha$ | AB4618 | S. Emr |
| *pRS313-BEM1* | Top10 | AB4762 | This study |
| *pRS313-bem1-Nterm* | Top10 | AB4682 | This study |
| *pRS313-bem1-Cterm* | Top10 | AB4652 | This study |
| *pRS313-Bem1-SH3aΔ* | Top10 | AB4653 | This study |
| *pRS313-Bem1-SH3bΔ* | Top10 | AB4655 | This study |
| *pRS313-Bem1-PXΔ* | Top10 | AB4654 | This study |

| | | | |
|---|---|---|---|
| *pRS313-Bem1-PB1Δ* | Top10 | AB4683 | This study |
| *pRS313-Bem1-SH3a-PB1* | Top10 | AB4745 | This study |
| *pRS313-Bem1-SH3b-PB1* | Top10 | AB4746 | This study |
| *pRS316-Bem1* | Top10 | AB4578 | This study |

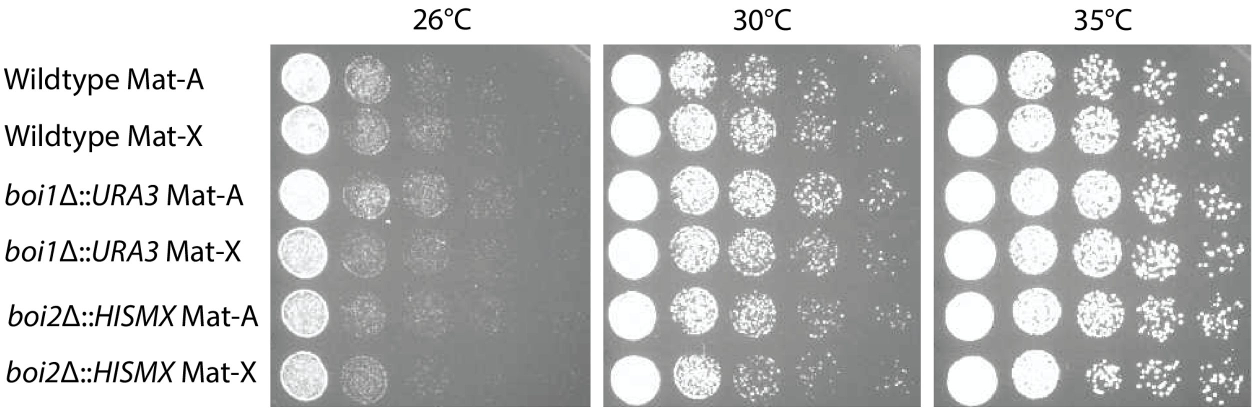


**Supplemental Figure 1.1:** *boi1*Δ or *boi2*Δ growth. 10-fold dilution assays of wildtype, *boi1Δ::URA3*, and *boi2Δ::HISMX* of each mating type at three temperatures: 26°C, 30°C, and 35°C on YPD for 24 hours.

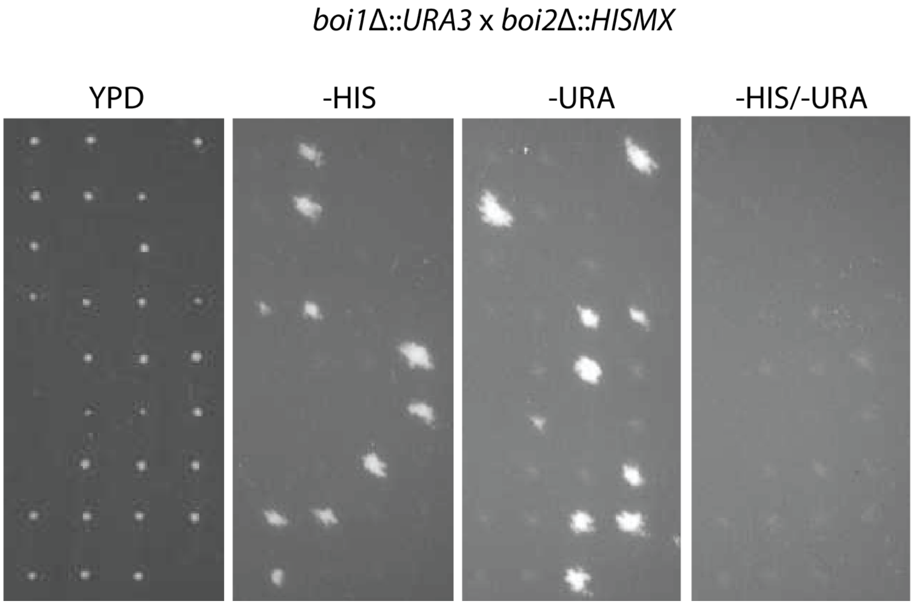


**Supplemental Figure 1.2:** Either Boi1 or Boi2 is essential for cell survival in the BY4741/2 background. Tetrad dissections of mated cells: *boi1Δ::URA3* x *boi2Δ::HISMX* diploid cells on YPD. Tetrads were replica plated on -HIS to select for *boi2Δ::HISMX*, -URA to select for *boi1Δ::URA3*, and -HIS/-URA to select for *boi1Δ::URA3 boi2Δ::HISMX*. There were no viable *boi1Δ::URA3 boi2Δ::HISMX* spores. All plates were grown for 2 days at 26°C.

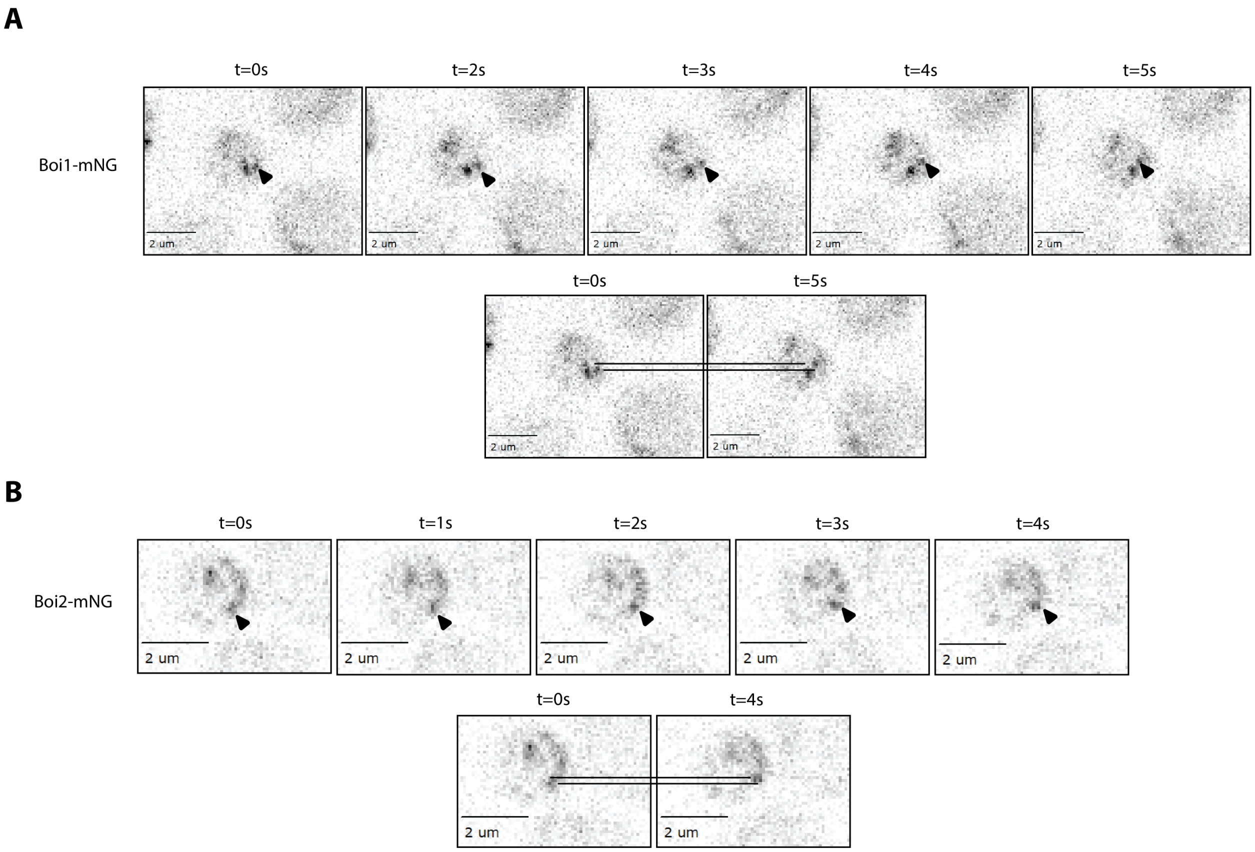


**Supplemental 2.1:** Lateral movement of Boi1-mNG and Boi2-mNG along the bud cortex. (A) Example of lateral patch movement in Boi1-mNG taken from videos similar to Video 1 with 50ms exposure timelapse captured every 160ms. Patch moves upwards along the cortex (highlighted with black arrows), slightly. Lower panel shows the displacement of the patch from the first timepoint to the last. (B) Example of lateral patch movement in Boi2-mNG taken with the same parameters as in A.

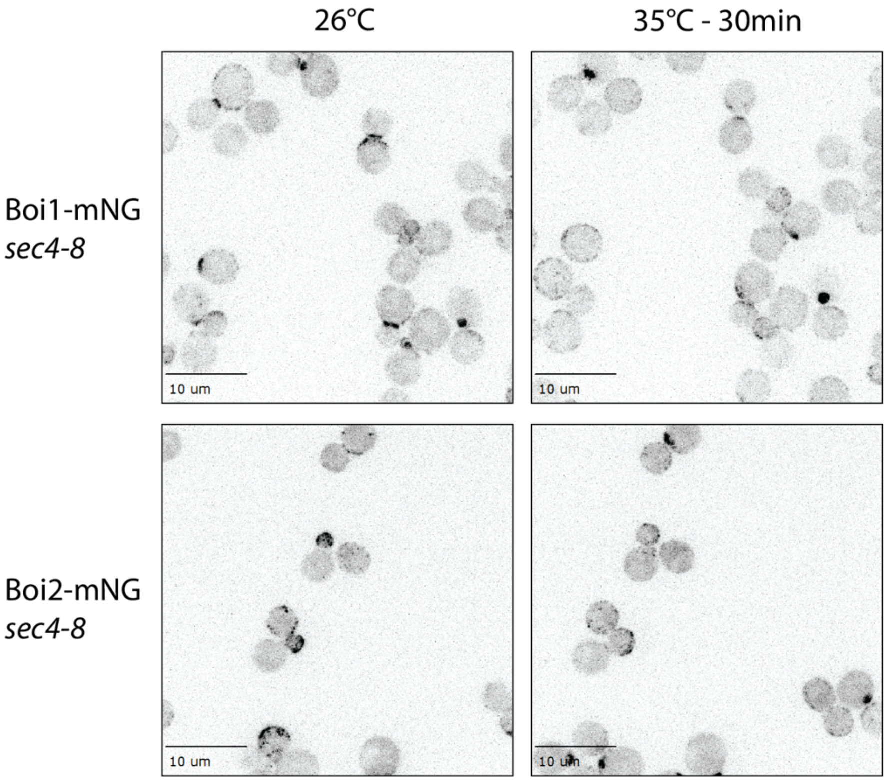


**Supplemental Figure 3.1:** Boi1-mNG and Boi2-mNG in *sec4-8* cells. Boi1-mNG and Boi2-mNG in *sec4-8* cells either at 26°C or shifted to the restrictive temperature, 35°C for 30 minutes on the CherryTemp. Cells were imaged with 15 plane z-stacks and were exposed for 200ms. Scale bars are 10µm.

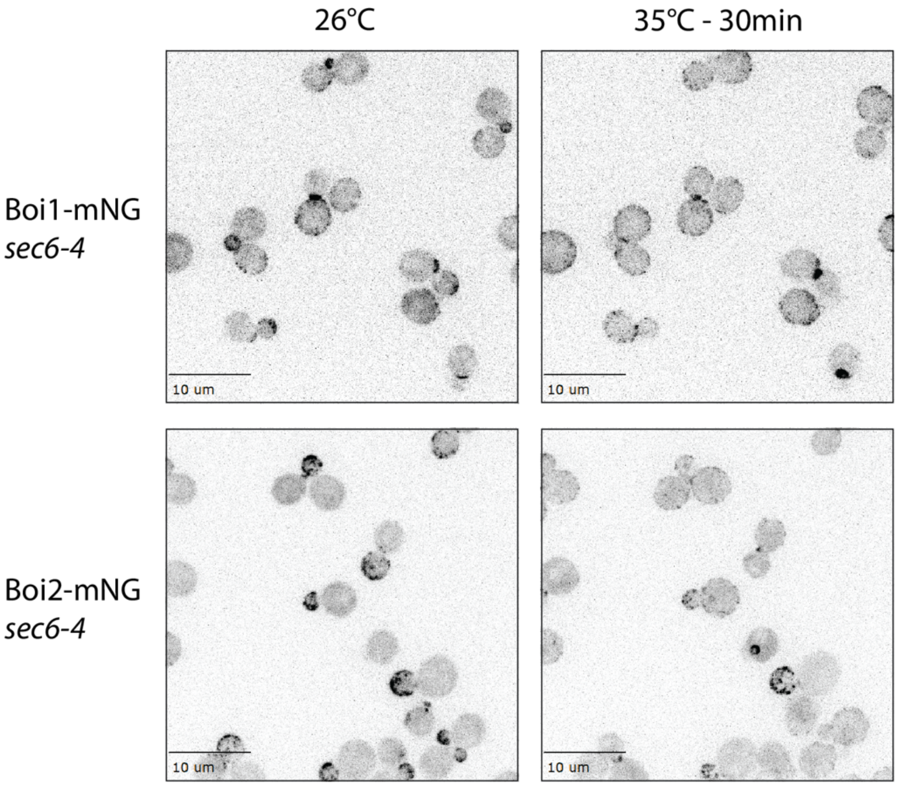


**Supplemental Figure 3.2:** Boi1-mNG and Boi2-mNG in *sec6-4* cells. Boi1-mNG and Boi2-mNG in *sec6-4* cells either at 26°C or shifted to the restrictive temperature, 35°C for 30 minutes on the CherryTemp. Cells were imaged with 15 plane z-stacks and were exposed for 200ms. Scale bars are 10µm.

Boi1-mNG *sec6-4* at 35°C - mid-cell single slice

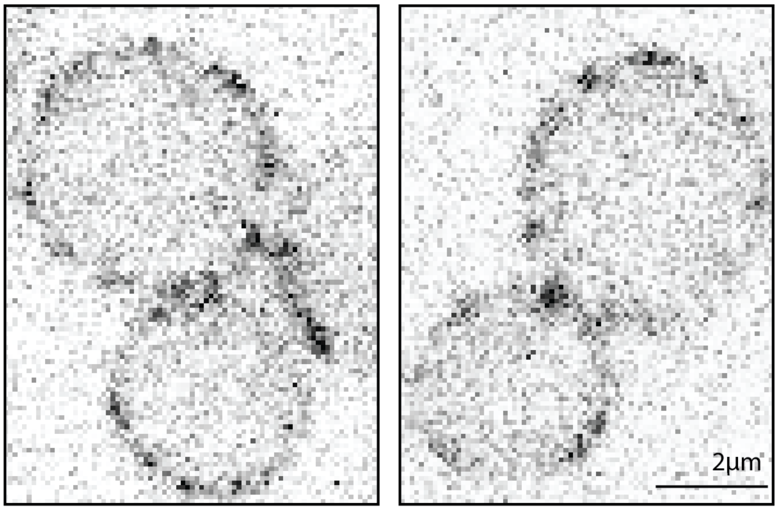


Boi2-mNG *sec6-4* at 35°C - mid-cell single slice

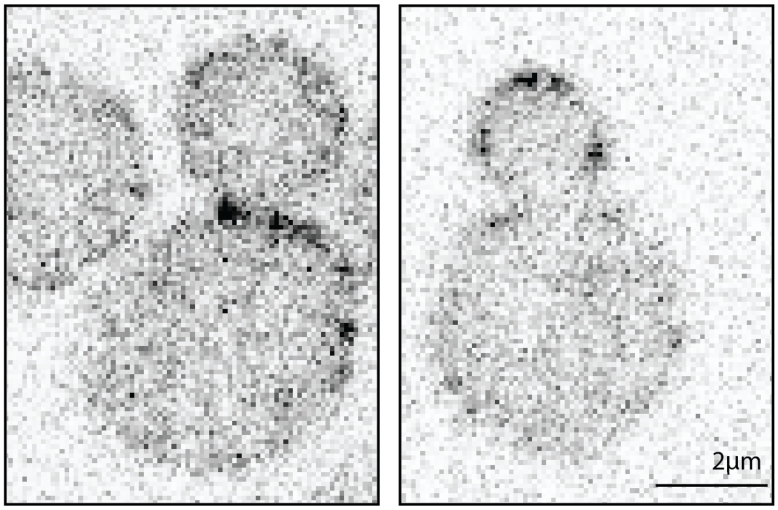


**Supplemental Figure 3.3:** Single slice images from Supplemental Figure 2.2 through the middle of the cell. Boi1-mNG and Boi2-mNG in *sec6-4* cells shifted to 35°C for over 30 minutes on the Cherry Temp. Exposure of 200ms. Scale bars are 2µm.

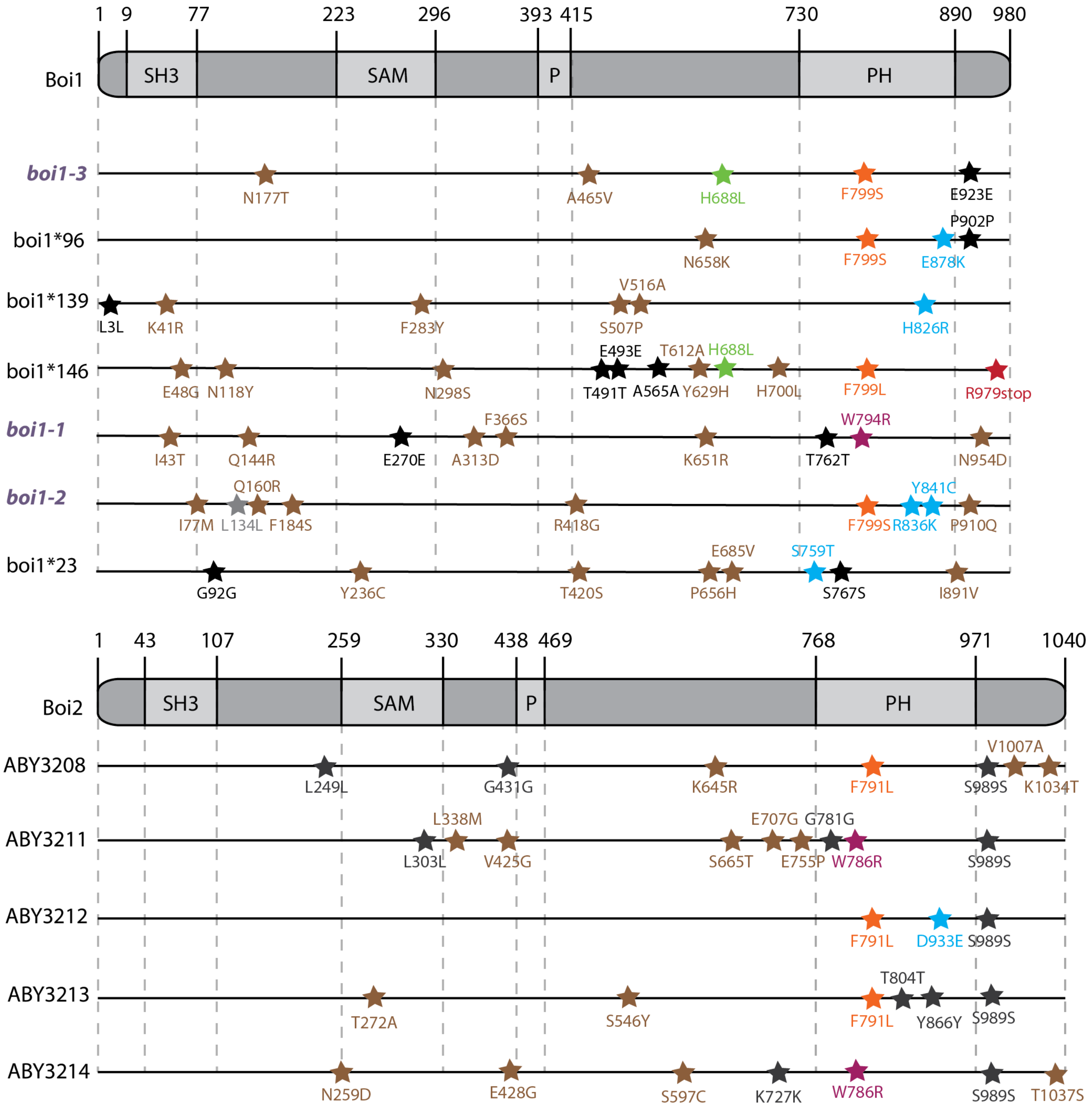


**Supplemental Figure 4.1:** Mutations in the temperature sensitive candidates in *BOI1* or *BOI2* mutants. Mutations found throughout *BOI1* and *BOI2* in several candidates along with the chosen temperature sensitive mutants (*boi1-1, boi1-2*, and *boi1-3*, highlighted in purple). Interesting PH domain mutations are highlighted in blue and green, along with missense mutations in brown and silent mutations in black to show the mutational coverage of the different screens.

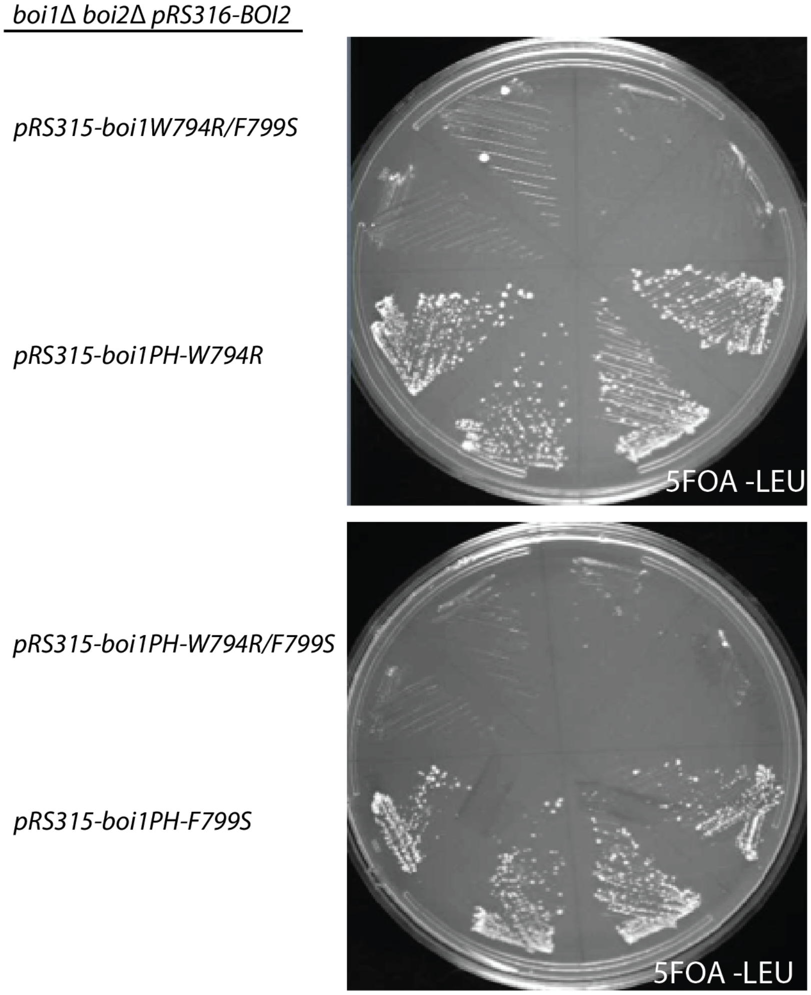


**Supplemental Figure 4.2:** Double mutants of *boi1W794R/F799S* are inviable. Plates showing four isolates with pRS315 plasmids carrying*: boi1W794R/F799S, boi1PH-W794R, boi1PH-W794R/F799S*, or *boi1PH-799S*. Plasmids were transformed into *boi1Δ boi2Δ pRS316-BOI2* cells and then plated on 5FOA -LEU to select against pRS316-BOI2 plasmid. Cells were grown for 3 days at 26°C.

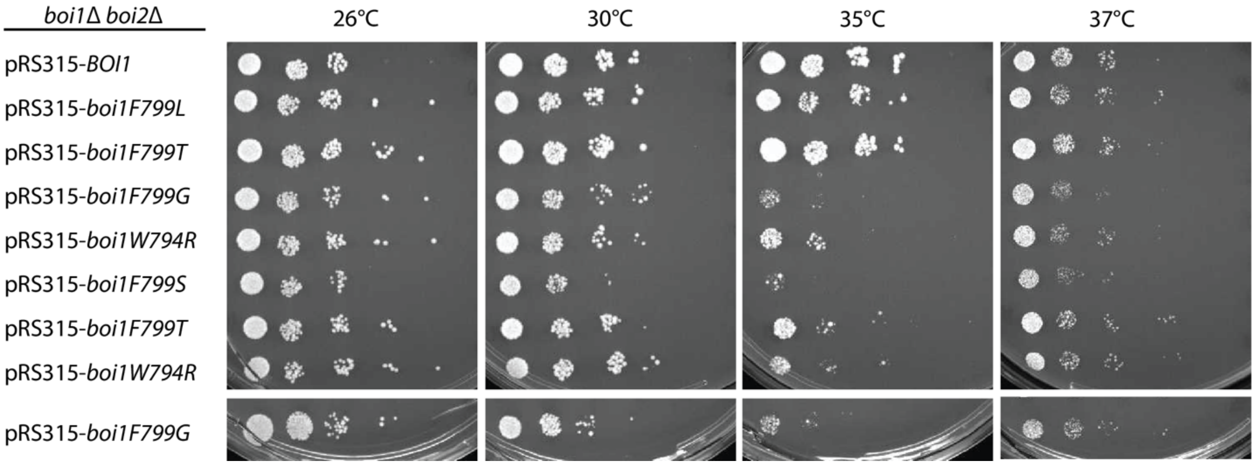


**Supplemental Figure 4.3:** Mini-screen of *Boi1W794X* and *boi1F799X* show that these positions are critical for function. Plates showing viable candidates from the screens looking at Boi1W794 and Boi1F799 mutated to virtually every other amino acid possible in yeast. The most sensitive candidates are: *Boi1W794R, Boi1W794G, Boi1F799S*, and *Boi1F799G*. Plates were grown at their respective temperatures for 2 days on SC -LEU.

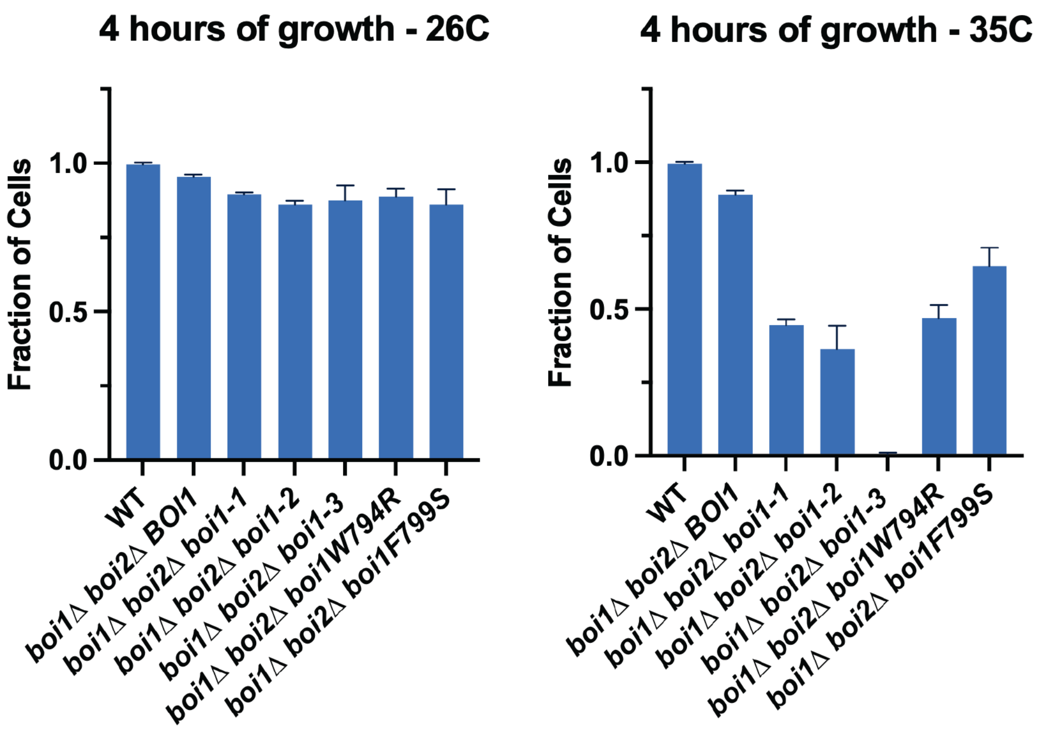


**Supplemental Figure 5.1:** Percentage of viable cells alive after incubation at respective temperatures. Data shows percentage of viable cells alive at the end of a 4-hour time-period at the respective temperatures. Data was collected by observing cells in at least 5 different fields of view with at minimum n=100. Graphs are a fraction of viable cells analyzed in each cell type.

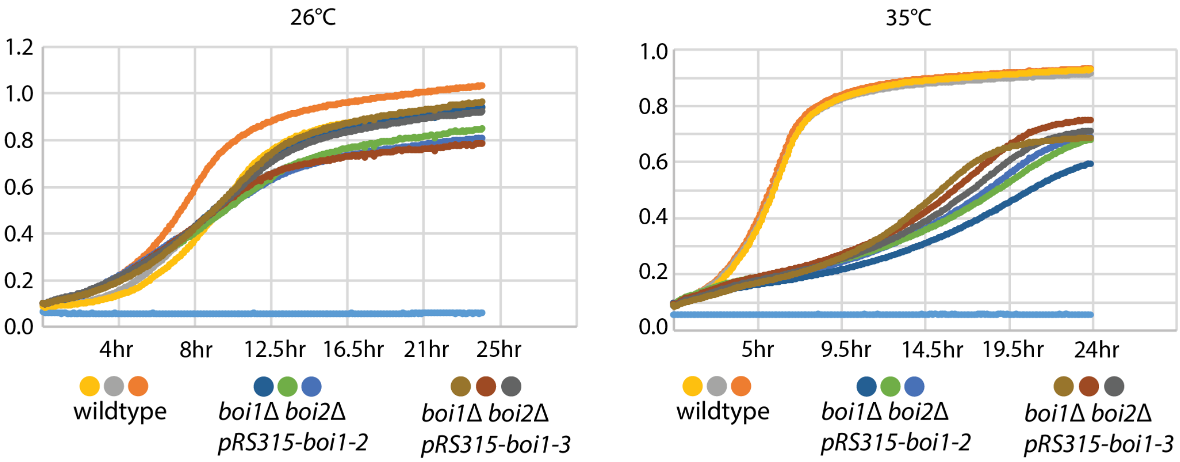


**Supplemental Figure 5.2:** Growth curves of *boi1* mutants compared to wildtype at permissive and restrictive temperatures. Graphs show optical density at 600nm of the cells grown at 26°C and at 35°C - three replicates of each cell type are shown including a blank (shown in cyan).

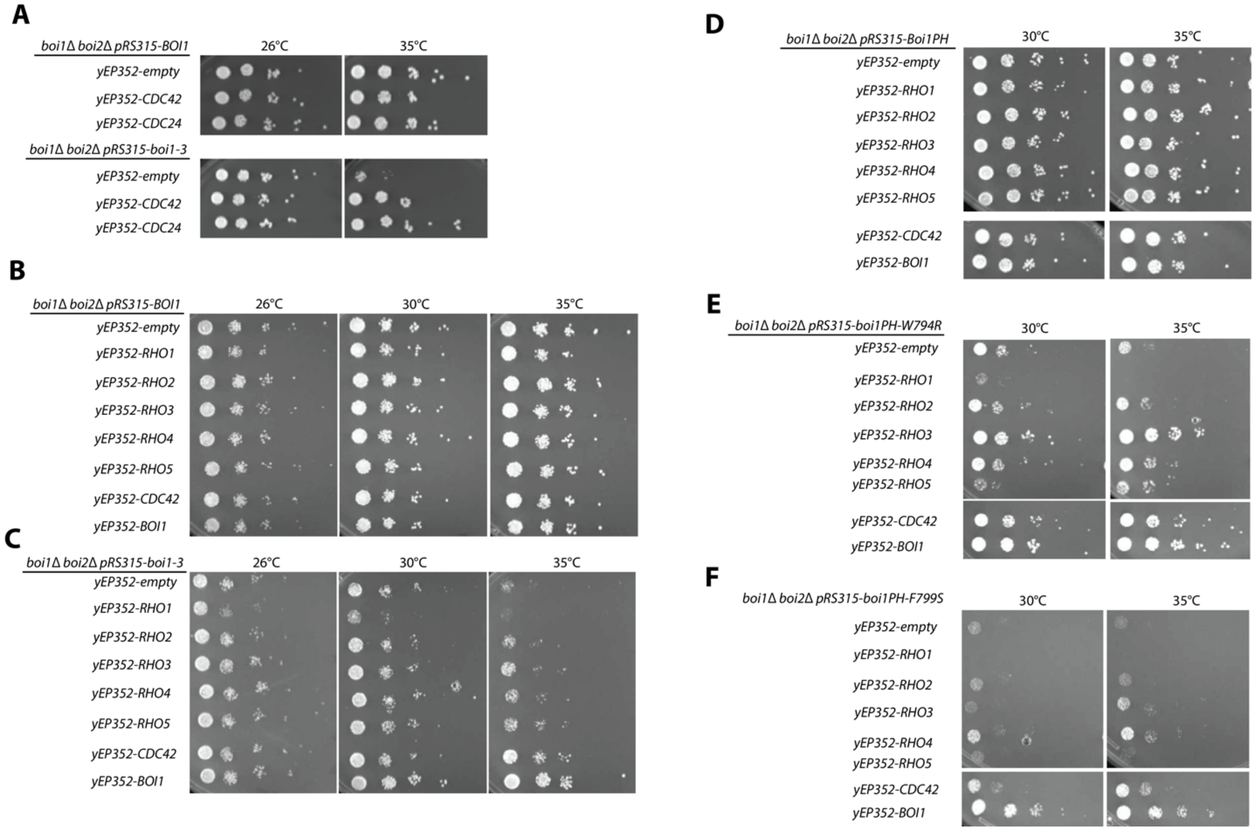


**Supplemental Figure 6.1:** Suppression of boi1 conditional mutants by over-expression of the Cdc42 pathway. (A) Growth assays of wildtype *BOI1* or *boi1-3* in a *boi1Δ boi2Δ* cells carrying yEP352 plasmids with either an empty plasmid, *CDC42*, or the GAP *CDC24*. Cells were grown on SC -URA/-LEU media for 2 days. (B and C) Growth assays of wildtype *BOI1* or *boi1-3* in a *boi1Δ boi2Δ* background with yEP352 plasmids either empty or with: *RHO1, RHO2, RHO3, RHO4, RHO5, CDC42*, or *BOI1*. Cells were grown on SC-URA/-LEU media for 2 days. (D, E, and F) Growth of *boi1Δ boi2Δ* background cells with a pRS315 vector containing Boi1PH (Boi1-730-980 C-terminal region) (D), *boi1PH-W794R* (E), or *boi1PH-F799S* (F), with the addition of yEP352 vector empty, or with the indicated Rho GTPases, similar to panels B and C. Cells were plated on SC -URA/-LEU and grown at respective temperatures for 2 days.

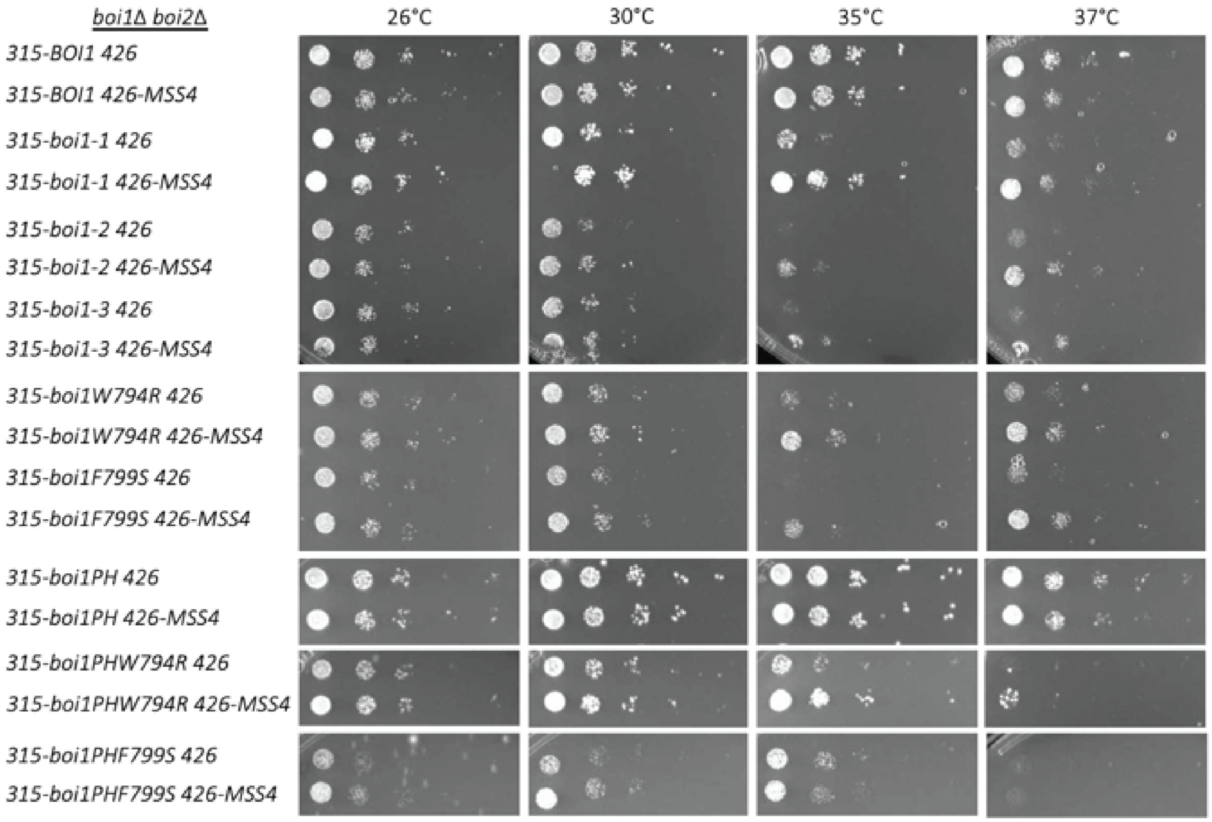


**Supplemental Figure 6.2:** Growth of *BOI1* mutants upon over-expression of *MSS4*. *MSS4* overexpressed with pRS426 2µm plasmid transformed into each of the indicated mutants. Dilution assays plated on -LEU/-URA plates and incubated at their respective temperatures for 2 days.

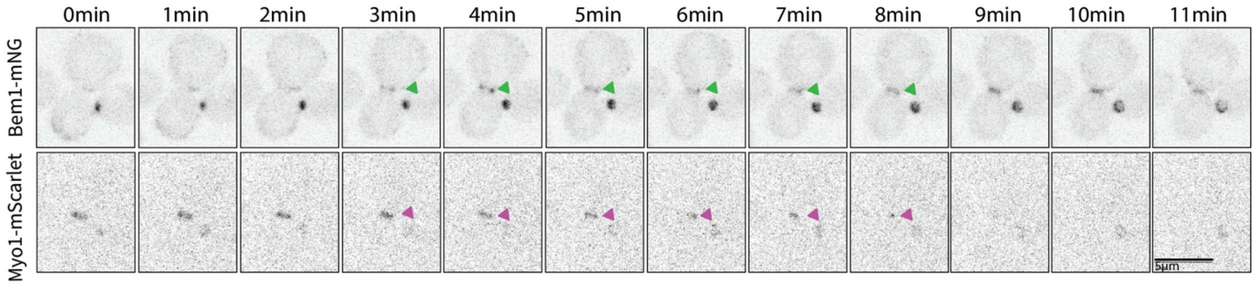


**Supplemental Figure 7.1:** Bem1-mNG colocalizes with Myo1-mScarlet prior to cytokinesis for 6 minutes. Still images of Bem1-mNG with Myo1-mScarlet. Images are max z-projections of 15 planes covering 4µm of the cell. Arbitrary timepoint "0min" is 3 minutes before Bem1-mNG's arrival to the bud neck. Scale bars are 5µm. See Figure 1E for Boi1-mNG compared to Myo1-mScarlet.

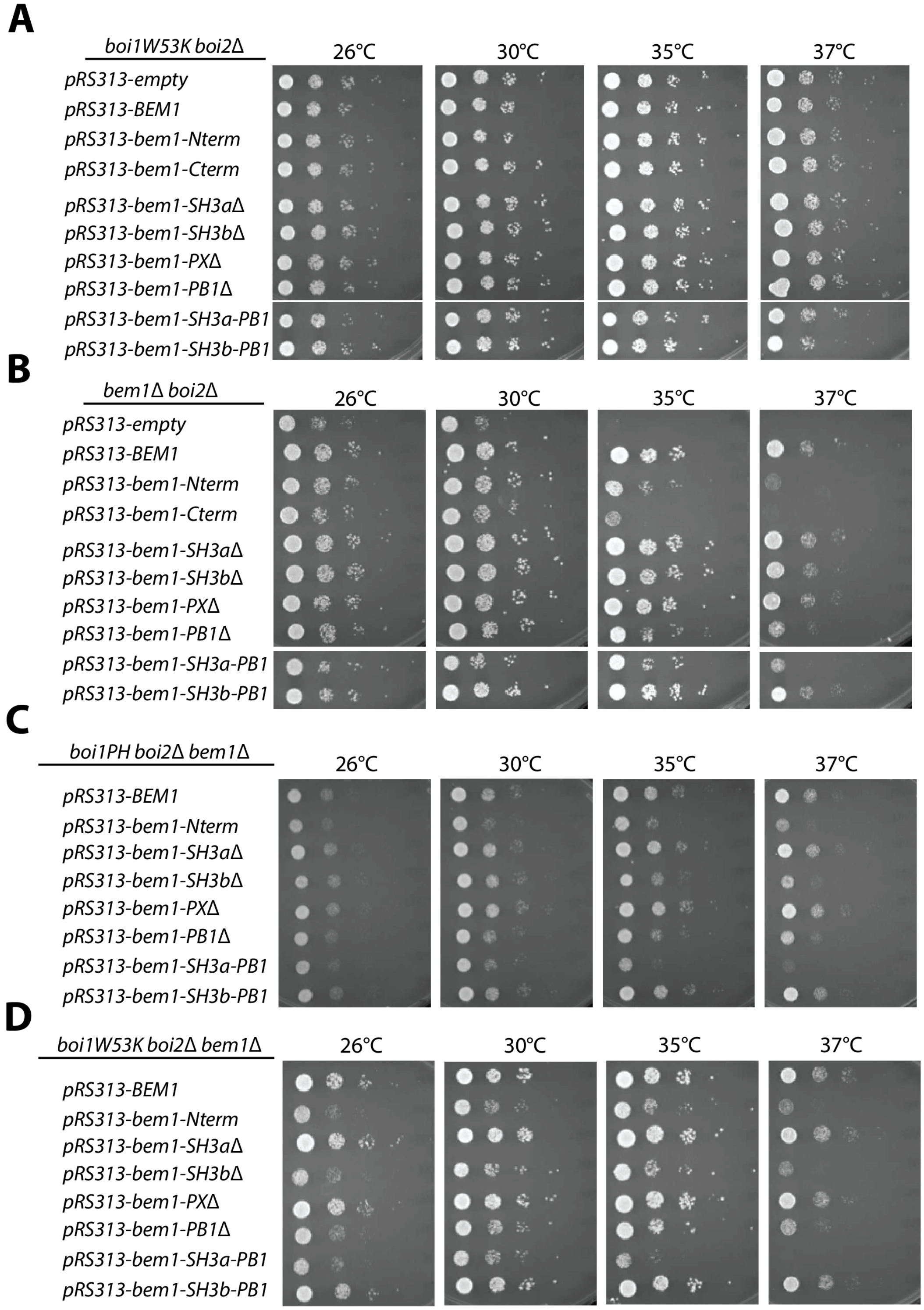


**Supplemental Figure 7.2:** Growth assays of Bem1 truncations in control strains and in strains where Boi1's SH3 domain is compromised. (A-D) Dilution assays of different Bem1p truncations in *boi1W53K boi2Δ BEM1* (A), *BOI1 boi2Δ bem1Δ* (B), *Boi1PH boi2Δ bem1Δ* (C), and *boi1W53K boi2Δ bem1Δ* (D). Cells were grown on -HIS plates for 3 days at the respective temperatures.

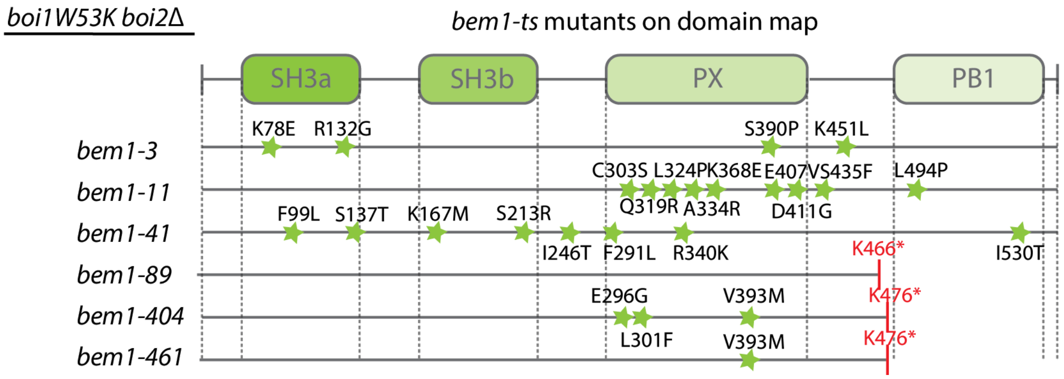


**Supplemental Figure 8.1:** Location of mutations in temperature sensitive candidates of *BEM1*. Termination codons are indicated in red by the preceding amino acid residue.

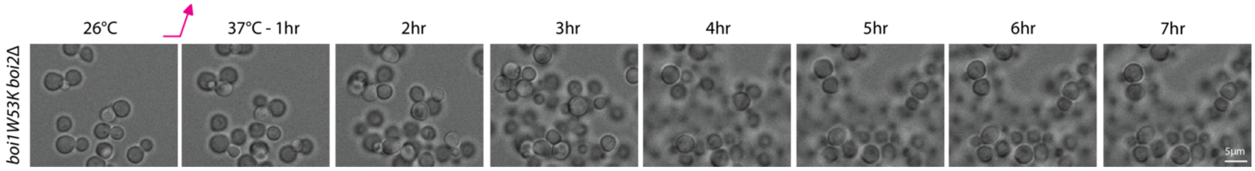


**Supplemental Figure 8.2:** DIC images of *boi1W53K boi2Δ* cells during growth. Cells were imaged once every five minutes for several hours after shifting to the restrictive temperature of 37°C. Scale bars are 5µm.
